# Histone H3K9 Methyltransferases Regulate Cortical Growth by Coordinating Heterochromatin Formation and Neural Progenitor Dynamics

**DOI:** 10.64898/2026.01.23.701405

**Authors:** Sophie Warren, Chris Hemmerich, Ram Podicheti, José-Manuel Baizabal

## Abstract

DNA packaging into heterochromatin is a fundamental mechanism of transcriptional silencing. However, how heterochromatin regulates neurogenesis in the developing cerebral cortex remains poorly understood. A defining feature of heterochromatin is trimethylation of histone H3 lysine 9 (H3K9me3), catalyzed by the H3K9 methyltransferases SETDB1, SUV39H1, and SUV39H2. Here, we generate a cortex-specific triple knockout mouse model lacking *Setdb1*, *Suv39h1*, and *Suv39h2* to interrogate the collective functions of H3K9 methyltransferases and H3K9me3 during corticogenesis. Loss of H3K9 methyltransferases disrupts cell-cycle dynamics and cortical neurogenesis, resulting in microcephaly. We show that H3K9me3 is associated with the silencing of distinct gene families, lineage-inappropriate genes, and transposable elements, and that its loss is accompanied by local chromatin opening and enhanced transcription factor occupancy. Our findings suggest that H3K9me3 regulates neurogenesis in part by silencing the growth-inhibitory gene *Cdkn1c* in intermediate progenitors. These results underscore the critical role of heterochromatin in the temporal control of neurogenesis.

## INTRODUCTION

The development of the mammalian cerebral cortex is orchestrated by neural stem cells (NSCs), also known as radial glia, which sequentially generate deep-layer neurons, upper-layer neurons, and glial cells^1^. NSCs either differentiate directly into cortical neurons or produce intermediate progenitor cells (IPCs), which typically undergo one or two rounds of cell division before terminal differentiation^1^. Precise regulation of NSC and IPC proliferation and cell cycle dynamics is essential to ensure an appropriate number of neurons and to establish the final size and laminar architecture of the cerebral cortex^1^.

Epigenetic mechanisms control gene expression via chemical modifications of chromatin. Among the best-characterized modifications are DNA methylation and histone methylation or acetylation, which collectively determine the degree of chromatin compaction and its accessibility to the transcriptional machinery^2^. Loosely packed chromatin, or euchromatin, permits transcription factor (TF) binding and is associated with active gene expression^2^. In contrast, tightly compacted chromatin, or heterochromatin, restricts TF access, leading to transcriptional repression^2^. Previous studies have identified multiple chromatin-modifying enzymes as critical regulators of cortical neurogenesis^3–7^. However, the role of heterochromatin-mediated gene silencing on the temporal progression of neurogenesis remains incompletely understood.

A distinctive feature of heterochromatin is trimethylation of histone H3 lysine 9 (H3K9me3), a repressive modification catalyzed by SETDB1, SUV39H1, and SUV39H2 (hereafter collectively referred to as H3K9 methyltransferases)^8^. H3K9me3 plays a critical role in maintaining genome stability by silencing repetitive genomic elements and has also been implicated in the repression of lineage-inappropriate genes during tissue development^9,10^. In the developing and adult mammalian brain, H3K9me3 displays cell-type-specific enrichment patterns, suggesting a role in neural fate specification and maintenance^11^. Consistent with this notion, individual H3K9 methyltransferases regulate embryonic and adult neurogenesis^4,12^. Additionally, deleterious mutations in *Setdb1* and *Suv39h2* have been associated with autism spectrum disorder and other neurodevelopmental syndromes^13,14^. However, loss of individual H3K9 methyltransferases results in only modest decreases in global H3K9me3 levels, likely reflecting functional redundancy among these enzymes^4,14,15^. Consequently, the mechanisms through which H3K9me3-marked heterochromatin regulates gene silencing and cortical neurogenesis remain unresolved.

Here, we generated a triple knockout (TKO) mouse line targeting *Setdb1*, *Suv39h1*, and *Suv39h2* to selectively deplete H3K9me3 in the embryonic cortex. Our data demonstrate that SETDB1, SUV39H1, and SUV39H2 are collectively required for cortical growth by regulating cell cycle dynamics, progenitor proliferation, neurogenesis, and cell survival. Integrative transcriptomic and epigenomic analyses revealed that H3K9me3 loss is associated with increased chromatin accessibility, elevated H3K27ac, enhanced TF occupancy, and aberrant gene transcription in the embryonic cortex. In addition, the loss of H3K9me3 is accompanied by activation of transposable elements (TEs), a process linked to upregulation of proximal genes. Lastly, our results suggest that H3K9me3 controls IPC amplification and differentiation by silencing *Cdkn1c*, a key determinant of cortical size. Together, these findings uncover previously unrecognized roles for H3K9me3-marked heterochromatin in regulating neural progenitor dynamics and cortical growth.

## RESULTS

### H3K9 methyltransferases determine cortical size

To investigate the coordinated functions of H3K9 methyltransferases during mouse cortical development, we used the *Emx1^Cre^* line to generate cortical-specific loss-of-function alleles for *Setdb1* and *Suv39h1* on a *Suv39h2*-null background^16,17^. The *Emx1^Cre^* line drives recombination of floxed alleles before the onset of cortical neurogenesis at embryonic day 9.5 (E9.5)^17^. We found that the *Suv39h1*/*Suv39h2* double knockout (DKO) cortex exhibited normal thickness and laminar organization at post-natal day 10 (P10) (Figure S1A). In contrast, the *Setdb1*/*Suv39h1* DKO cortex carrying a single wild-type copy of *Suv39h2* displayed a pronounced reduction in cortical thickness (Figure S1A). Microcephaly was even more severe in the TKO cortex, which lacked all functional alleles of *Setdb1*, *Suv39h1*, and *Suv39h2* (Figure S1A). In the TKO cortex, deep layers were disproportionately reduced compared to upper layers (Figure S1A).

Most cortical neurons in the TKO cortex displayed strong H3K9me3 depletion at P10 (Figure S1B), suggesting that SETDB1, SUV39H1, and SUV39H2 produce most H3K9me3 in the cortex, consistent with the low expression of *Setdb2*^18^. We also observed a few cells retaining normal H3K9me3 levels in the TKO cortex, likely representing cortical interneurons, which are not targeted by the *Emx1^Cre^* line (Figure S1B)^17^. Importantly, H3K9me2, another repressive mark, was largely unchanged in TKO neurons (Figure S1C), suggesting specific loss of H3K9me3.

We next examined the developmental time course of microcephaly in the TKO cortex. A slight reduction in cortical thickness was apparent at E15.5, becoming progressively more pronounced by E17.5 and P0 (Figure 1A). Quantification of cell number within a cortical column at P0 revealed a substantial decrease in the TKO, whereas cell density remained comparable to controls (Figure S2A). To assess temporal H3K9me3 depletion, we focused on the ventricular zone (VZ), the niche of cortical NSCs, and observed marked H3K9me3 loss at E15.5 and E17.5 (Figure 1B). Together, these results indicate that SETDB1, SUV39H1, and SUV39H2 establish H3K9me3 in the developing cortex and influence cortical growth.

**Figure 1.**
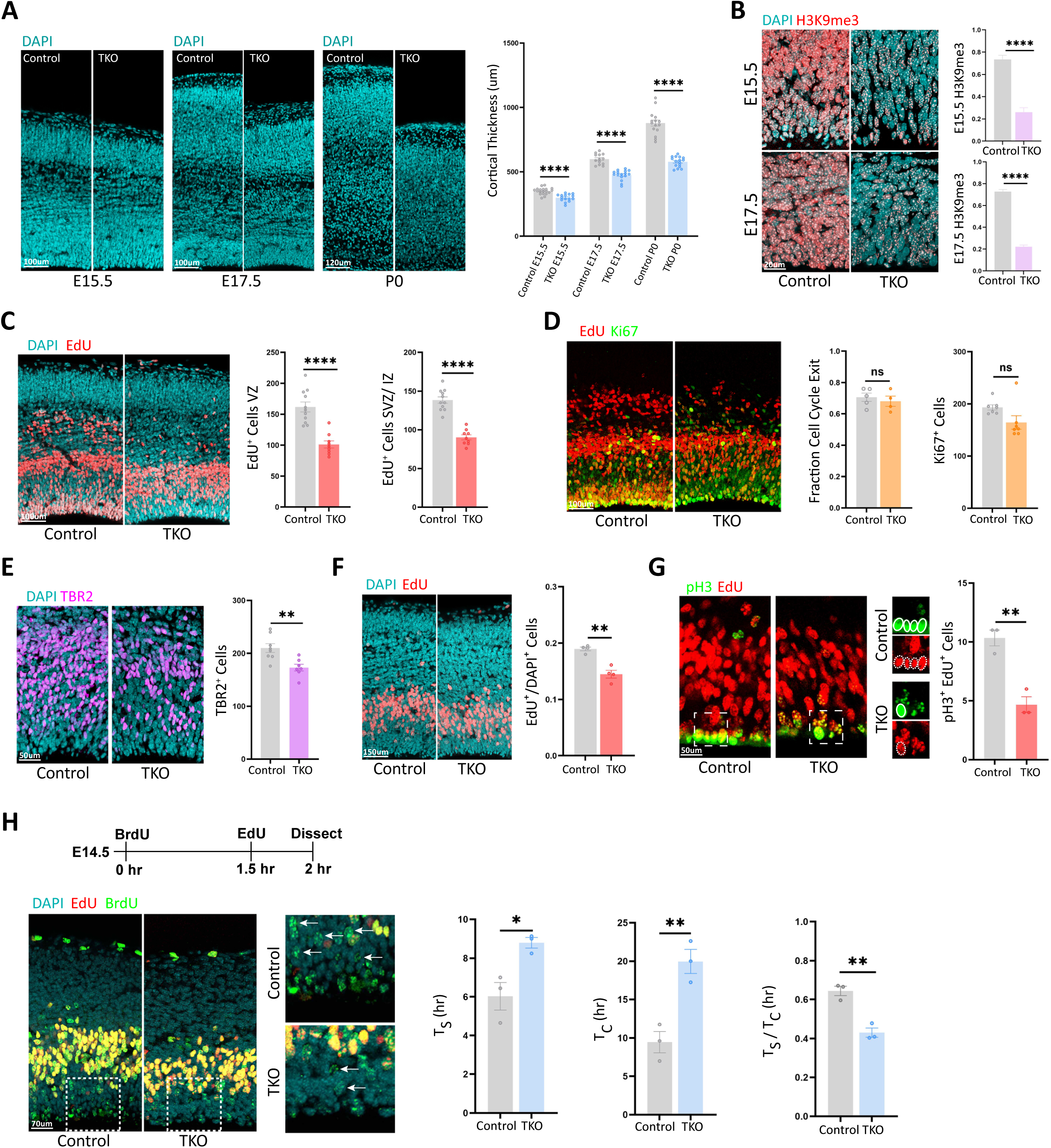
H3K9 methyltransferases regulate cell cycle dynamics and cortical growth. (A) Coronal sections of the E15.5, E17.5, and P0 developing cortex revealed a progressive reduction in cortical thickness in TKO mice. (B) H3K9me3 immunostaining of the VZ in E15.5 and E17.5 control and TKO cortices. Quantification of normalized pixel intensity confirmed a robust depletion of H3K9me3 in the TKO cortex. (C) EdU administration at E14.5 and analysis ∼24 h later showed an abnormal distribution of EdU⁺ cells in the TKO cortex, accompanied by a significant reduction of EdU⁺ cells in the VZ and SVZ/IZ. (D) Cell cycle exit analysis using EdU and Ki67 at E15.5 revealed no differences between genotypes. The number of proliferating neural progenitors (Ki67⁺ cells) was comparable in control and TKO cortices. (E) Reduced numbers of TBR2⁺ cells in the E15.5 TKO cortex. (F) EdU administration at E14.5, with analysis 1 hr. later, showed a decreased fraction of S-phase cells (EdU^+^/DAPI^+^ cells) in the TKO cortex. (G) EdU administration at E14.5, with analysis 3 hrs. later, revealed a significant reduction in labeled mitotic cells (pH3⁺ EdU⁺ cells) in the TKO cortex. (H) Schematic of BrdU and EdU administration and analysis. Cells labeled with both BrdU and EdU at the time of analysis (yellow cells) represent S-phase cells. Insets show higher-magnification examples of BrdU⁺ EdU⁻ cells (green cells, arrows), indicating that fewer TKO progenitors exited S-phase before EdU administration. Quantification of S-phase length (Ts) and total cell cycle length (Tc) showed an increase in TKO progenitors, whereas the relative S-phase duration (Ts/Tc) was reduced. Quantifications are shown as mean ± SEM. Unpaired Student’s *t* test: \**p* < 0.05, \*\**p* < 0.01, \*\*\*\**p* < 0.0001; ns, not significant.

### H3K9 methyltransferases regulate cell cycle dynamics during mid-neurogenesis

To determine whether the microcephaly phenotype arises from defects in cell proliferation and differentiation, we injected the thymidine analog 5-Ethynyl-2’-deoxyuridine (EdU) at E14.5 and analyzed EdU^+^ cell distribution approximately one day later (E15.5) in control and TKO cortices. EdU is incorporated into replicating DNA during S-phase, permanently labeling proliferating cells and their progeny. We observed a substantial reduction of EdU^+^ cells in the VZ, subventricular zone (SVZ), and intermediate zone (IZ) of the TKO cortex (Figure 1C). To assess whether this altered EdU distribution reflected changes in proliferation and differentiation, we measured cell cycle exit by quantifying the fraction of EdU^+^ cells negative for the proliferation marker Ki67 (MKI67). Surprisingly, the fraction of cells exiting the cell cycle was comparable between control and TKO cortices (Figure 1D), and the total number of proliferating Ki67^+^ cells was unchanged at E15.5 (Figure 1D). Furthermore, while the number of Pax6^+^ NSCs was normal, Tbr2^+^ IPCs were reduced (Figures 1E and S2B), suggesting that the TKO primarily affects IPC generation rather than NSC maintenance.

To investigate whether the altered EdU distribution reflected changes in cell cycle dynamics, we analyzed the proportion of cells in S-phase and mitosis at 1 and 3 hrs. post-EdU injection, respectively. The TKO cortex showed a significant reduction in both S-phase (EdU^+^) and mitotic (pH3^+^) cells (Figures 1F and 1G). Using a double-labeling paradigm with EdU and bromodeoxyuridine (BrdU)^19^, we measured cell cycle length and found a marked increase in both S-phase duration (Ts) and total cell cycle length (Tc) in the TKO cortex (Figure 1H). Although the S-phase was prolonged, it represented a smaller fraction of the total cell cycle (Ts/Tc; Figure 1H), suggesting that lengthening of other cell cycle phases contributes disproportionately to the overall increase in cycle duration. Collectively, these findings indicate that H3K9 methyltransferases are critical for regulating cell cycle dynamics and IPC production during mid-neurogenesis in the developing cortex.

### H3K9 methyltransferases control proliferation, differentiation, and survival during late neurogenesis

To address the impact of *Setdb1*, *Suv39h1*, and *Suv39h2* loss on later cortical development, we injected EdU at E16.5 and analyzed EdU^+^ cells approximately one day later (E17.5). Like our observations at E14.5, the number of EdU^+^ cells was strongly reduced in the VZ, SVZ, and IZ of the TKO cortex (Figure 2A). However, unlike mid-neurogenesis, the E17.5 TKO cortex exhibited a significant increase in the fraction of cells exiting the cell cycle, indicative of enhanced differentiation at this stage (Figure 2B). This was consistent with a decrease in Ki67^+^ cells (Figure 2C). Analysis of progenitor populations revealed that the number of PAX6^+^ cells remained unchanged, whereas TBR2^+^ cells increased at E17.5 (Figure 2D, S2C). Moreover, many TBR2^+^ cells were Ki67^−^ in the TKO cortex (Figure 2E), suggesting that a larger fraction of IPCs had differentiated into immature cortical neurons. To assess whether neuronal fate specification was altered, we labeled neurons with EdU at E13.5 and examined their laminar distribution at P5 (Figure S2D). The proportion of EdU^+^ neurons in upper cortical layers tended to increase in the TKO cortex, although this change did not reach statistical significance (Figure S2D), indicating that neuronal fate specification was largely preserved.

**Figure 2.**
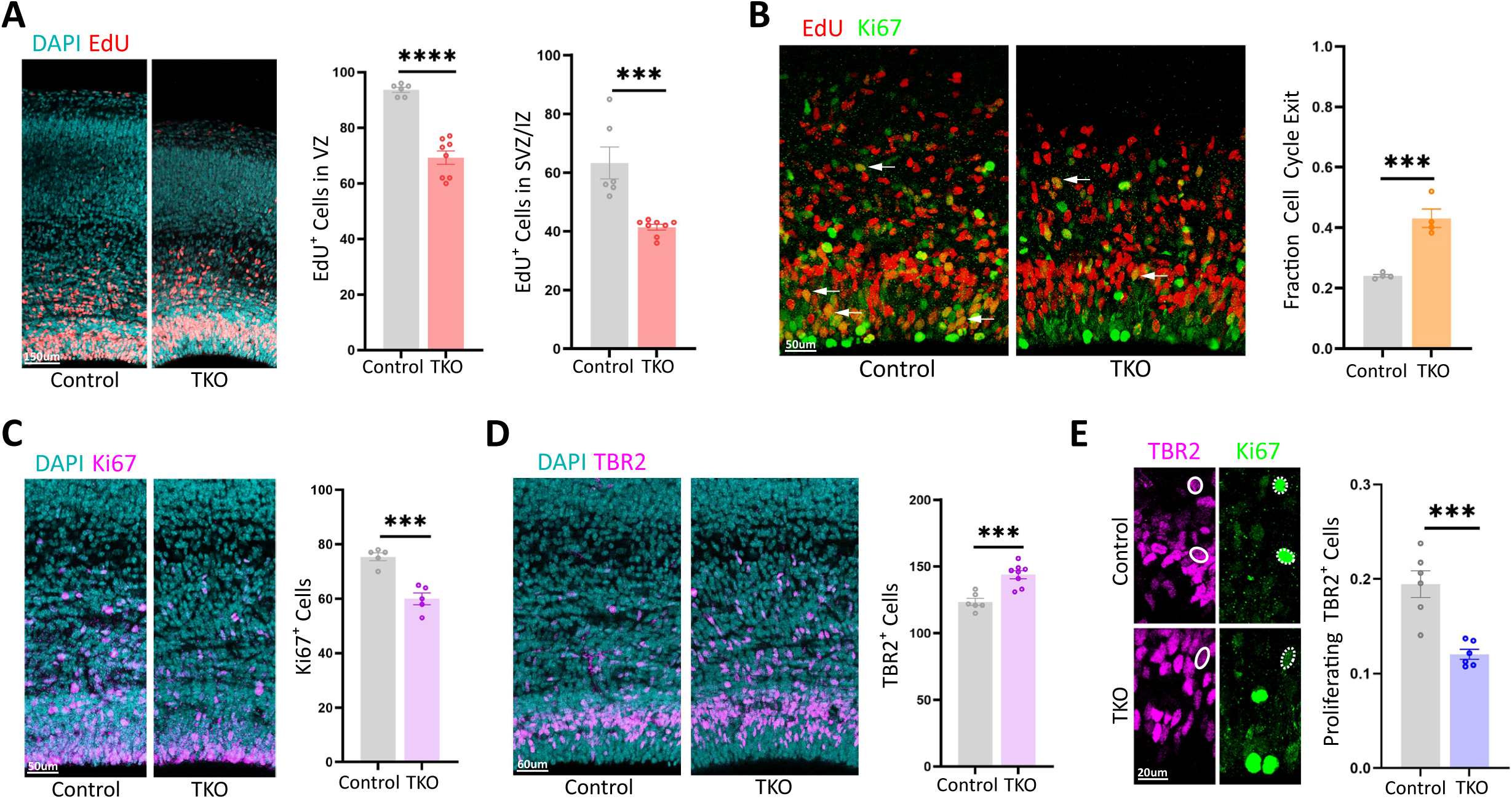
H3K9 methyltransferases regulate neural progenitor proliferation and neurogenesis. (A) EdU administration at E16.5 with analysis ∼24 hrs. later (E17.5) revealed an atypical distribution of EdU⁺ cells in the TKO cortex, accompanied by a reduction in EdU⁺ cells in the VZ and SVZ/IZ. (B) A reduced number of cells re-entering the cell cycle (EdU⁺ Ki67⁺ cells, arrows indicate some examples) was observed in the E17.5 TKO cortex ∼24 hrs. after EdU administration. Accordingly, the fraction of cells exiting the cell cycle was significantly increased in the TKO cortex. (C) The number of proliferating neural progenitors (Ki67⁺ cells) decreased in the E17.5 TKO cortex. (D) The number of TBR2⁺ cells increased in the E17.5 TKO cortex. (E) The fraction of TBR2⁺ Ki67⁺ cells (dashed circles) relative to the total number of TBR2⁺ cells was reduced in the E17.5 TKO cortex. Quantifications are shown as mean ± SEM. Unpaired Student’s *t* test: \*\*\**p* < 0.001.

We next examined the role of H3K9 methyltransferases in neuronal survival by assessing cleaved caspase 3 (CC3). While no increase in apoptosis was observed at E15.5, the number of CC3^+^ cells was elevated in the E17.5 and P0 TKO cortex, predominantly within the cortical plate, indicating impaired neuronal survival (Figure S2E). To determine whether this cell death was linked to genome instability, we analyzed γH2A.X, a marker of DNA double-strand breaks, but found no obvious increase in TKO neurons (Figure S2F), suggesting that DNA damage is not the primary driver of apoptosis. Together, these results demonstrate that H3K9 methyltransferases determine cortical growth by regulating neural progenitor proliferation, differentiation, and neuronal survival during late neurogenesis.

### H3K9 methyltransferases silence cell adhesion and lineage-inappropriate genes in the embryonic cortex

To identify genes regulated by H3K9 methyltransferases during cortical development, we performed bulk RNA-seq at E15.5, a stage at which the TKO cortex displays mild microcephaly but no detectable cell death. Principal component analysis revealed a clear separation between control and TKO transcriptomes (Figure S3A). Differential expression analysis identified 446 downregulated and 643 upregulated genes in the TKO cortex (Figure 3A; Table S1). To resolve cell-type-specific transcriptional changes, we next performed single-cell RNA-seq (scRNA-seq) at E14.5. Clustering analysis identified 15 transcriptionally distinct cell populations in the control cortex (Figure 3B). Importantly, corresponding clusters were preserved in the TKO cortex (Figure 3C), indicating that overall cell-type identities remained largely intact despite transcriptional perturbations. Approximately 50% of the differentially expressed genes (DEGs) identified by bulk RNA-seq were also detected in the scRNA-seq dataset (Figure S3B), demonstrating substantial concordance between the two approaches.

**Figure 3.**
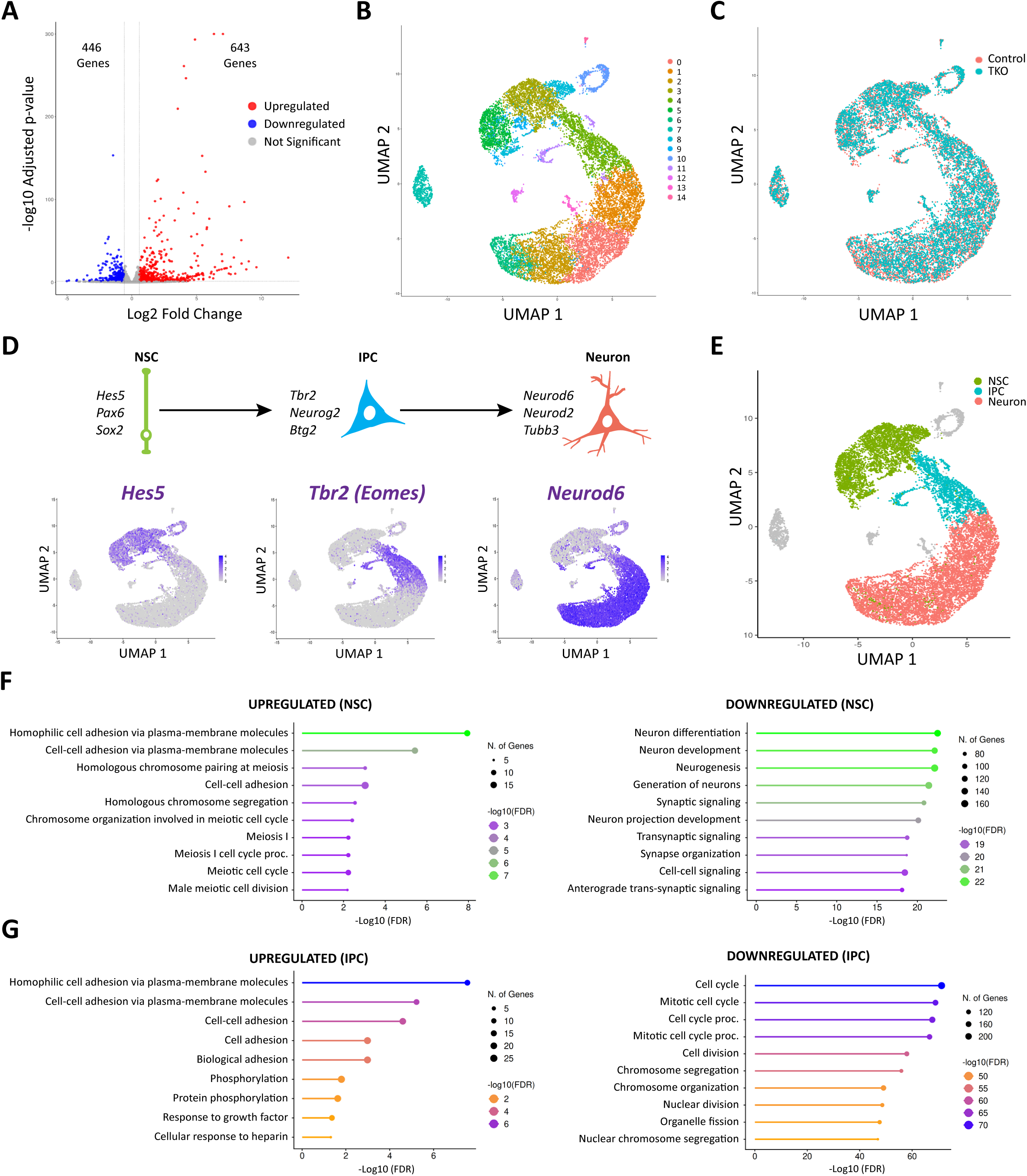
H3K9 methyltransferases silence cell adhesion and lineage-inappropriate genes in the embryonic cortex. (A) Volcano plot showing DEGs identified by bulk RNA-seq comparing E15.5 control and TKO cortices. The number of upregulated and downregulated genes in the TKO cortex are indicated. (B) UMAP plot showing 15 distinct transcriptional clusters identified by scRNA-seq in E14.5 control cortices. (C) Overlap of control and TKO scRNA-seq datasets demonstrates preservation of overall transcriptional identities in the TKO cortex. (D) Schematic illustrating the three major stages of cortical lineage differentiation: NSCs, IPCs, and cortical neurons, with representative marker genes indicated. UMAP plots highlight cells with high expression of *Hes5*, *Tbr2* (*Eomes*), and *Neurod6*. (E) UMAP plot delineating NSCs, IPCs, and neurons within the cortical lineage based on expression of the predefined markers shown in (D). (F) Gene ontology analysis of DEGs between control and TKO NSCs revealed enrichment of upregulated categories related to cell adhesion and meiosis, and downregulation of genes associated with neuronal differentiation. (G) Gene ontology analysis of DEGs between control and TKO IPCs showed upregulation of cell adhesion pathways and downregulation of genes involved in cell cycle progression and proliferation.

Using established marker genes, we identified a continuum of cortical lineage progression from NSCs (*Hes5*, *Pax6*, *Sox2*), to IPCs (*Tbr2*, *Neurog2*, *Btg2*), and projection neurons (*Neurod6*, *Neurod2*, *Tubb3*) (Figures 3D, 3E, and S3C–S3E)^20^. This approach enabled the identification of cell-type-specific DEGs within the TKO cortex (Table S1). Gene ontology analysis of upregulated genes revealed a shared enrichment for cell–cell adhesion molecules across NSCs, IPCs, and neurons (Figures 3F, 3G, and S3F). Notably, meiosis-associated genes, which are normally restricted to germ cells and silenced in the brain, were aberrantly upregulated specifically in TKO NSCs (Figure 3F; Table S1). In contrast, downregulated genes in NSCs were enriched for functions related to neurogenesis and neuronal differentiation (Figure 3F). In IPCs, downregulated transcripts were selectively associated with cell cycle progression and mitotic division (Figure 3G). Finally, in cortical neurons, downregulated genes were enriched for pathways involved in synaptic function (Figure S3G), suggesting impaired or delayed neuronal maturation. Together, these results demonstrate that loss of H3K9 methyltransferases leads to transcriptional dysregulation across the cortical lineage, affecting neural progenitor identity, cell cycle control, and neuronal development.

### H3K9me3 loss is associated with enhanced chromatin accessibility in the developing cortex

To map genomic regions marked by H3K9me3 in the E14.5 control cortex, we performed chromatin immunoprecipitation and sequencing (ChIP-seq). Spike-in chromatin was used to normalize ChIP efficiency, and only uniquely mapped reads were analyzed in both control and TKO samples. Irreproducible Discovery Rate (IDR) analysis of scaled ChIP-seq data identified 51,053 highly reproducible H3K9me3 peaks in control cortices (Figure S4A). In contrast, only 1,200 reproducible peaks were detected in the TKO cortex (Figure S4B), indicating a near-complete loss of H3K9me3. Consistently, visualization of H3K9me3 across the genome confirmed a strong reduction in TKO samples (Figure 4A).

**Figure 4.**
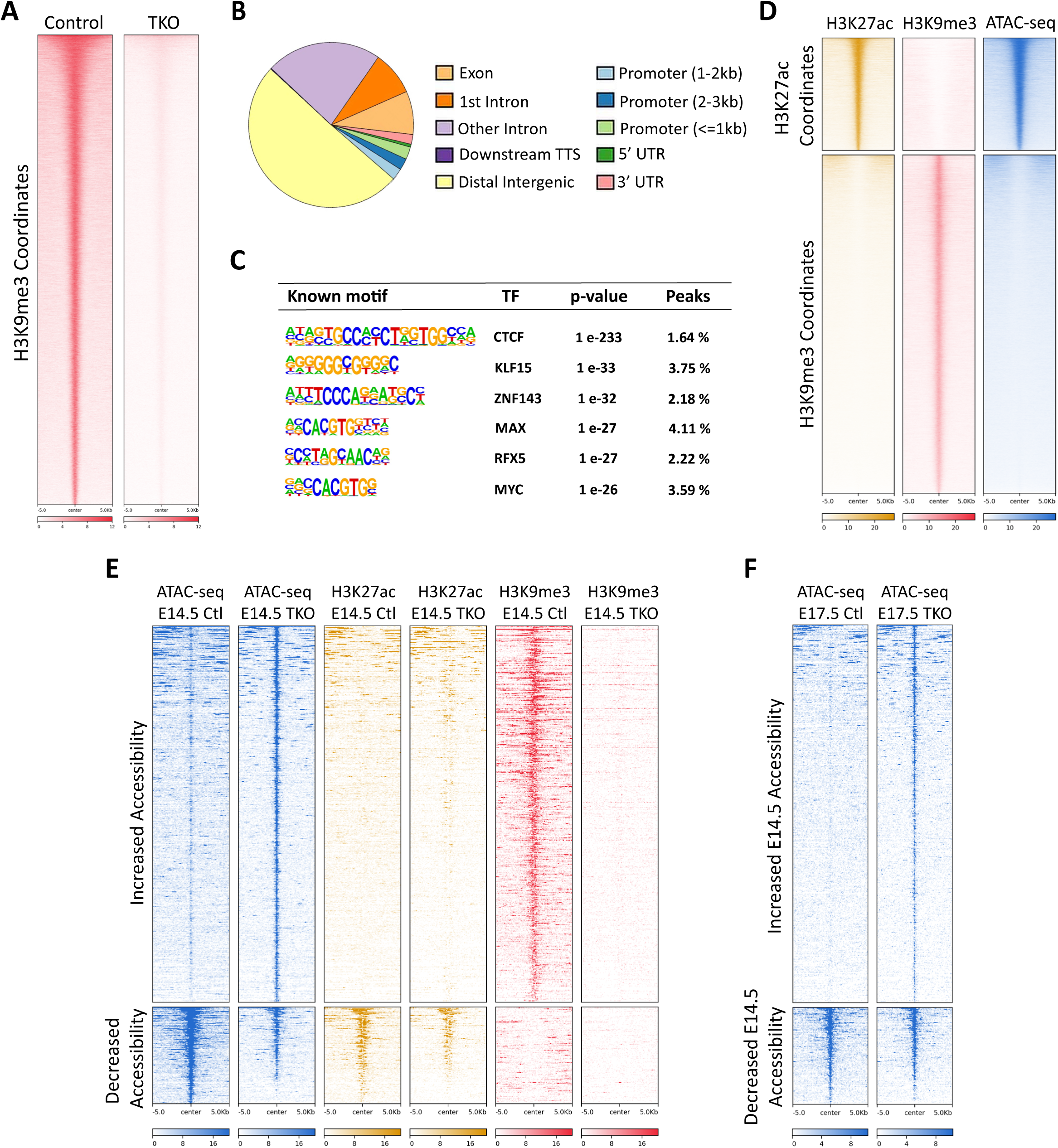
H3K9me3 is associated with heterochromatin formation in the embryonic cortex. (A) Heatmaps showing H3K9me3-enriched genomic regions in E14.5 control cortices and their depletion in TKO cortices. (B) Pie chart depicting the genomic distribution of H3K9me3 in E14.5 cortical cells, with the majority of signal localized to intergenic and intronic regions. (C) DNA motif enrichment analysis within H3K9me3 ChIP-seq peaks from E14.5 control cortex. The top six enriched TF motifs are shown. (D) Heatmaps displaying H3K27ac, H3K9me3, and ATAC-seq signals in E14.5 control cortex. H3K9me3 predominantly decorated genomic regions characterized by low H3K27ac and low chromatin accessibility. (E) Heatmaps of ATAC-seq, H3K27ac, and H3K9me3 signals at a subset of chromatin regions in E14.5 control (Ctl) and TKO cortices. Regions that gained accessibility in TKO cortices were marked by H3K9me3 in controls, and some acquired moderate levels of H3K27ac. Regions that lost accessibility in TKO cortices were not associated with H3K9me3 loss. (F) Heatmaps showing ATAC-seq signal at E17.5 over chromatin regions that gained or lost accessibility at E14.5, as defined in (E). Regions that increased accessibility at E14.5 remained open at E17.5 in the TKO cortex.

In control cortices, H3K9me3 peaks were predominantly localized to intergenic and intronic regions, with a smaller fraction mapping to promoters and exons (Figure 4B). Motif analysis of H3K9me3-decorated regions revealed a significant enrichment of binding motifs for CTCF, KLF5, ZNF143, MAX, RFX5, and MYC (Figure 4C). The top enrichment of CTCF motifs is consistent with recent reports demonstrating a role for H3K9me3 in preventing CTCF binding and regulating chromatin looping^21,22^. H3K9me3 was largely excluded from regions enriched for H3K27ac, a marker of active promoters and enhancers (Figure 4D)^23^. Moreover, H3K9me3-marked regions exhibited low chromatin accessibility, as assessed by the Assay for Transposase-Accessible Chromatin with high-throughput sequencing (ATAC-seq) (Figure 4D). Together, these data indicate that H3K9me3 primarily marks heterochromatin in the developing cortex.

To examine whether H3K9me3 loss impacts chromatin accessibility, we conducted ATAC-seq in E14.5 control and TKO cortices. We identified 572 regions with differential accessibility between genotypes (Figure S4C), of which 445 regions gained and 117 lost accessibility in the TKO cortex (Figure 4E and S4C). The majority of the differentially accessible regions were located within intergenic and intronic sequences (Figure S4D). To assess the persistence of these changes, we performed ATAC-seq at E17.5, when proliferation and differentiation defects are prominent in the TKO cortex (Figure 2B and 2C). Regions that gained accessibility at E14.5 remained highly accessible at E17.5 (Figure 4F), indicating lasting alterations in chromatin compaction following H3K9me3 loss.

We next examined whether altered chromatin accessibility was accompanied by changes in histone acetylation. Spike-in-normalized H3K27ac ChIP-seq identified 168 regions with increased and 63 regions with decreased H3K27ac in the E14.5 TKO cortex (Figure S4E). Most differentially acetylated regions were located at promoters (Figure S4F). In addition, the sequences that gained accessibility in the TKO cortex exhibited moderate-to-marginal increases in H3K27ac (Figure 4E). Importantly, genomic regions that gained accessibility in the TKO cortex were decorated by H3K9me3 in control cortices, whereas regions that lost accessibility lacked H3K9me3 in controls (Figure 4E). Collectively, these findings demonstrate that H3K9me3 loss is associated with increased chromatin accessibility at a subset of loci, supporting a direct role for H3K9 methyltransferases in maintaining chromatin compaction during cortical development.

### H3K9me3 loss is associated with activation of specific gene families and lineage-inappropriate genes

To identify genes potentially silenced by H3K9me3 in the embryonic cortex, we mapped H3K9me3 to annotated promoters. We identified 1,536 genes with H3K9me3 occupancy within ±1.5 kb of transcription start sites (TSS) (Table S2). Most of these promoters were also enriched for H3K4me3 (Figure 5A), indicating a transcriptionally active or poised state. At these promoters, H3K9me3 typically flanked the central H3K4me3 peak (Figures 5B and 5C). Among the 1,536 putative gene targets of H3K9me3, 100 were upregulated, and 33 were downregulated in the TKO cortex (Figure 5D; Table S2). Importantly, the upregulated genes exhibited larger expression fold changes than the downregulated genes (Figure 5D; Table S2), consistent with a primary role for H3K9me3 in transcriptional repression.

**Figure 5.**
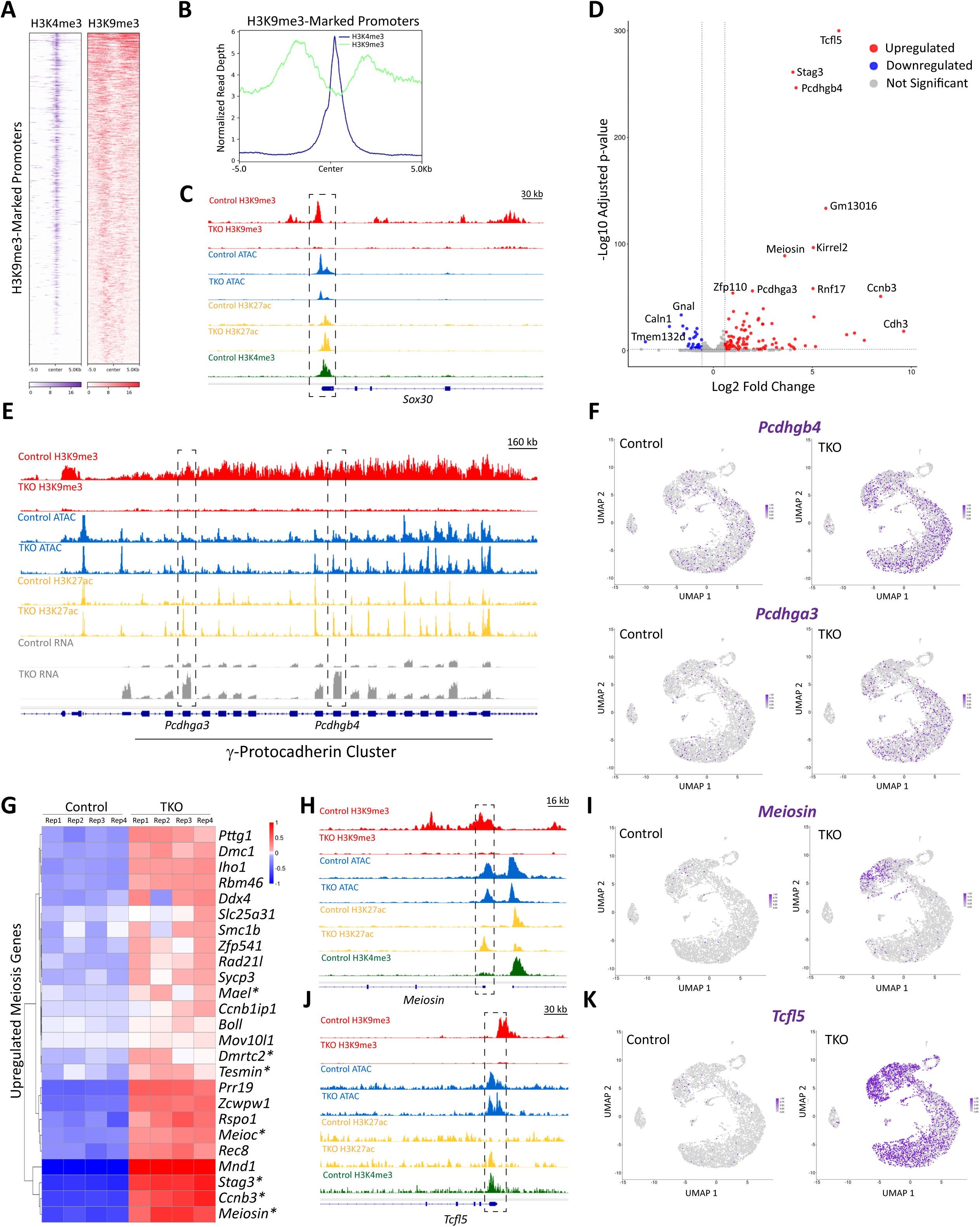
H3K9me3 is associated with repression of protocadherin clusters and meiosis genes in the embryonic cortex. (A) Heatmap showing enrichment of H3K9me3 at a subset of annotated promoters in the E14.5 control cortex. Most of these promoters were also marked by H3K4me3, indicating an active or poised transcriptional state. (B) Aggregate projection profiles of H3K9me3 and H3K4me3 across the genomic regions shown in (A) revealed that maximal H3K9me3 enrichment typically flanked the H3K4me3 peak at promoters. (C) Genome browser tracks illustrating partial overlap between H3K9me3 and H3K4me3 at the *Sox30* promoter (dashed rectangle) in the E14.5 control cortex. (D) Volcano plot of DEGs in the TKO cortex that displayed H3K9me3 at their promoters in the WT cortex. The most strongly dysregulated genes are labeled, including *Pcdhgb4*, *Pcdhga3*, *Meiosin*, and *Tcfl5*. (E) Genome browser tracks showing a broad H3K9me3 domain spanning the γ-protocadherin gene cluster. Loss of H3K9me3 in the TKO cortex was accompanied by moderate increases in chromatin accessibility, H3K27ac enrichment, and upregulation of *Pcdhga3* and *Pcdhgb4* (dashed rectangles). (F) UMAP plots from scRNA-seq data showed sparse expression of *Pcdhga3* and *Pcdhgb4* in the control cortex and their upregulation across the cortical lineage in the TKO cortex. (G) Heatmap of meiotic genes upregulated in the TKO cortex based on bulk RNA-seq data. Asterisks denote meiotic genes marked by H3K9me3 at their promoters in the WT cortex. (H) Genome browser tracks showing H3K9me3 enrichment at the *Meiosin* promoter. Low H3K4me3 levels correlated with the absence of *Meiosin* expression in the control E14.5 cortex. Loss of H3K9me3 in the TKO cortex was associated with increased H3K27ac at the *Meiosin* promoter (dashed rectangle). (I) scRNA-seq data indicated specific upregulation of *Meiosin* in NSCs of the E14.5 TKO cortex. (J) Genome browser tracks showing H3K9me3 enrichment at the *Tcfl5* promoter. Depletion of H3K9me3 in the TKO cortex coincided with increased chromatin accessibility and H3K27ac at this locus (dashed rectangle). (K) scRNA-seq data showed widespread transcriptional activation of *Tcfl5* in the E14.5 TKO cortex.

A prominent class of upregulated H3K9me3 target genes encodes cell–cell adhesion molecules, including *Cdh3* and multiple clustered protocadherin genes such as *Pcdhgb4* and *Pcdhga3* (Figure 5D; Table S2). Clustered protocadherin genes are organized within large genomic regions, and recent evidence indicates that they play a role in suppressing neural progenitor proliferation by antagonizing Wnt signaling^24,25^. In the developing cortex, the alpha, beta, and gamma protocadherin clusters were decorated by broad H3K9me3 domains (Figures 5E, S5A, and S5B). In total, 35 protocadherin genes were upregulated and 6 were downregulated in the E15.5 TKO cortex (Figure S5C). Loss of H3K9me3 across the gamma protocadherin cluster was associated with moderate, localized increases in chromatin accessibility and H3K27ac in the TKO cortex (Figure 5E). Single-cell analysis revealed that upregulated protocadherin genes exhibited a “salt-and-pepper” expression pattern across NSCs, IPCs, and cortical neurons (Figure 5F). Beyond protocadherins, we observed substantial H3K9me3 enrichment across additional gene families, including clusters encoding zinc finger proteins (*Zfp* genes) (Figure S5D), some of which were transcriptionally activated in the TKO cortex (Figure 5D; Table S2). Hence, broad H3K9me3 domains are associated with the silencing of a subset of gene families in the embryonic cortex.

Strikingly, we also identified aberrant activation of meiosis-associated genes, which are normally silent in the embryonic cortex (Figure 5G). A subset of these genes—including *Meiosin*, *Meioc*, and *Stag3*—displayed prominent H3K9me3 enrichment at their promoters and were predicted to be direct targets of H3K9me3-mediated repression (Figures 5H, S5E, and S5G). Consistent with our gene ontology analyses (Figures 3F and 3G), these genes were preferentially upregulated in NSCs of the TKO cortex (Figures 5I, S5F, and S5H). In addition, *Tcfl5*, a gene encoding a transcriptional regulator involved in spermatogenesis^26,27^, emerged as one of the strongest upregulated targets (Figure 5D). *Tcfl5* showed promoter-associated H3K9me3 in controls and was robustly upregulated across most cells in the TKO cortex (Figures 5J and 5K). In addition, *Tcfl5* and *Meiosin* activation upon H3K9me3 loss was accompanied by *de novo* accumulation of H3K27ac at their promoters (Figures 5H and 5J), reflecting a transition from silenced to active chromatin. These findings suggest that H3K9me3 participates in transcriptional repression of meiotic genes, thereby safeguarding neural identity during cortical development.

### H3K9me3 heterochromatin represses transposable elements and restricts transcription factor occupancy

A substantial fraction of H3K9me3 peaks in the embryonic cortex overlapped repetitive genomic elements (Table S3). To determine whether H3K9me3 is associated with repression of these elements, we focused on the 455 genomic regions that simultaneously lost H3K9me3 and gained chromatin accessibility in the TKO cortex (Figures 4E and 4F). These regions were highly enriched for TEs, including long terminal repeats (LTRs), long interspersed nuclear elements (LINEs), and short interspersed nuclear elements (SINEs) (Figure 6A; Table S3). Consistent with increased chromatin accessibility in the TKO cortex, bulk RNA-seq revealed activation of repetitive elements, including endogenous retroviruses (ERV1, ERVK, ERVL), DNA transposons (DNA, hAT-Blackjack, hAT-Tip100, TcMar-Tc2, TcMar-Tigger, UCON), retrotransposons (CR1, Gypsy), LTRs, LINEs (L1), SINEs (srpRNAs), and simple repeats (Figure 6B). Thus, loss of H3K9me3 is accompanied by both increased accessibility and transcriptional activation of TEs at a subset of genomic loci in the embryonic cortex.

**Figure 6.**
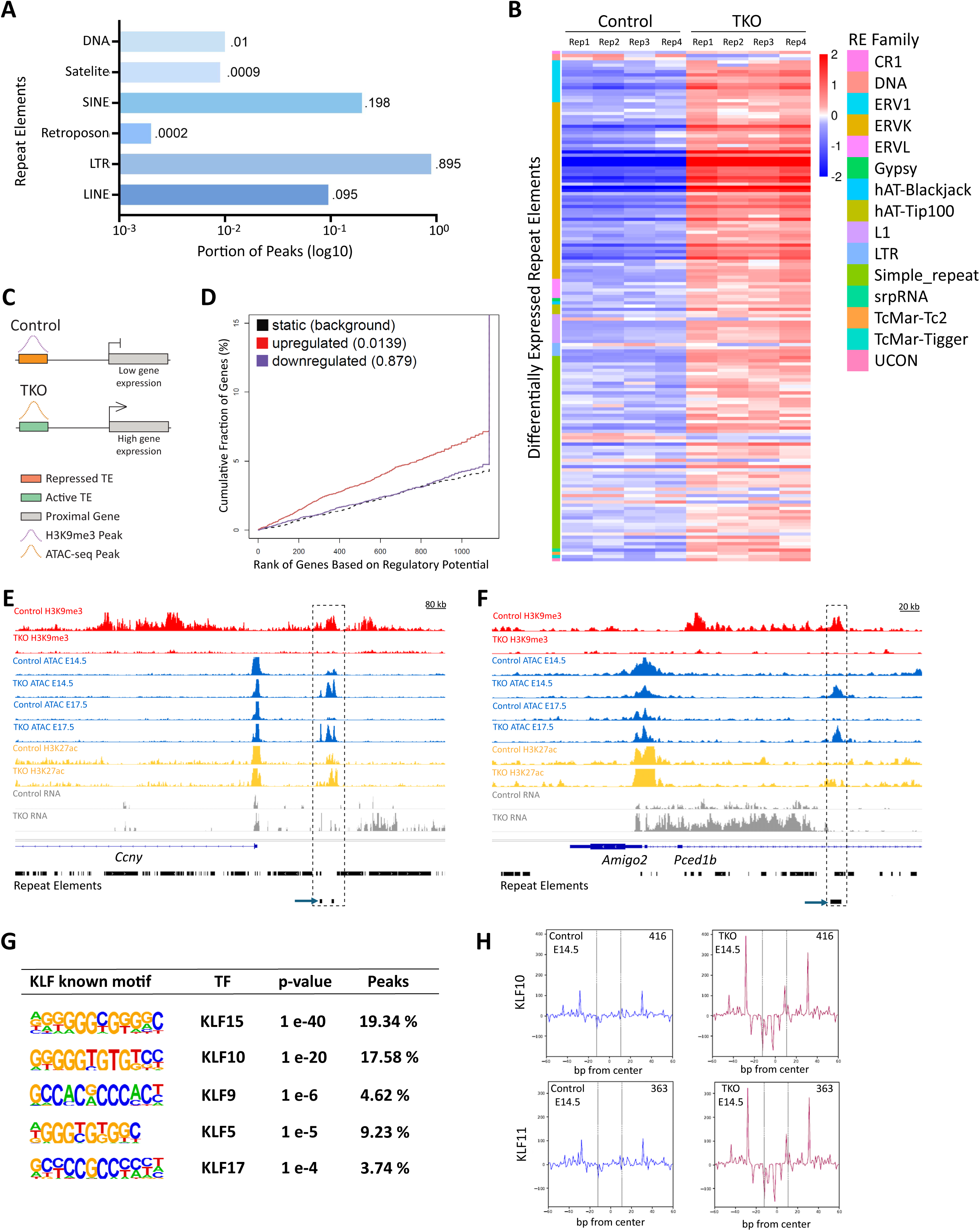
H3K9me3-marked heterochromatin represses transposable elements and restricts transcription factor occupancy. (A) Fraction of repeat element classes marked by H3K9me3 in the E14.5 control cortex that gained chromatin accessibility in the TKO cortex. (B) Bulk RNA-seq analysis revealed activation of repeat elements in the TKO cortex. The heatmap shows color-coded classification of dysregulated repeat element (RE) families (see main text for details). (C) Schematic illustrating TE activation and its association with upregulation of proximal genes in the TKO cortex. (D) BETA analysis demonstrated a significant association between TE activation and upregulation of proximal genes, whereas no significant association was observed for downregulated genes. The plot displays p-values and cumulative fractions of upregulated and downregulated genes relative to the background. (E) Genome browser tracks showing an H3K9me3-marked region upstream of *Ccny* that gained accessibility at E14.5 and E17.5 in the TKO cortex (dashed rectangle). This region contained a TE (IAPLTR2b), indicated by the arrow. RNA-seq tracks show concurrent upregulation of both IAPLTR2b and *Ccny* in the TKO cortex. (F) Genome browser tracks showing loss of H3K9me3 and increased accessibility within an intronic region of *Pced1b* (dashed rectangle). The arrow indicates the region with increased accessibility that partially overlapped with a simple repeat ([TCC]n) and a TE (SINE PB1D10) (Repeat Elements track). Loss of a broad H3K9me3 domain within the *Pced1b* intronic sequences was accompanied by robust transcriptional activation (RNA-seq tracks) of several TEs (Repeat Elements track) and *Pced1b* in the TKO cortex. (G) DNA motif enrichment analysis of loci that lost H3K9me3 and gained accessibility in the TKO cortex. Motifs for several KLF regulators were significantly enriched within these regions. (H) ATAC-seq footprinting analysis using the regions with increased chromatin accessibility in the TKO cortex as input. Enhanced footprint depth (shown between dashed lines) was observed for KLF10 and KLF11, indicating increased TF occupancy upon chromatin decompaction. The number of loci exhibiting increased KLF10 and KLF11 footprints is indicated. The regions flanking the footprints showed globally increased accessibility in the TKO cortex.

Active TEs can be co-opted as cryptic enhancers to regulate gene expression^28,29^. To test whether activated TEs were associated with gene upregulation in the TKO cortex, we applied Binding and Expression Target Analysis (BETA)^30^, which integrated genomic regions of increased chromatin accessibility with dysregulated genes within 100 kb (Figure 6C). To specifically examine distal regulatory elements, we excluded promoters (defined as ±1.5 kb around TSS), yielding 201 distal genomic regions—predominantly composed of repetitive elements—that formed 370 interactions with 351 dysregulated genes (Table S4). Notably, all 351 genes associated with these distal TEs were upregulated in the TKO cortex (Figure 6D; Table S4). Representative examples included the upregulated genes *Ccny* and *Pced1b*, which were linked to distal TEs that were actively transcribed in the TKO cortex (Figures 6E and 6F). These TEs also exhibited increased chromatin accessibility and elevated H3K27ac in the TKO cortex (Figures 6E and 6F). Together, our results reveal a link between TE activation and gene induction, supporting a role for TEs as potential cryptic enhancers in the TKO cortex.

To identify transcriptional regulators that may drive activation of promoters and TEs in the TKO cortex, we conducted DNA motif enrichment analysis on regions that lost H3K9me3 and gained chromatin accessibility. This analysis revealed strong enrichment for Krüppel-like factor (KLF) motifs (Figure 6G). To test whether reduced chromatin compaction in the TKO cortex facilitated KLF binding, we performed ATAC-seq footprinting analysis on the 455 affected regions^31^. We detected pronounced footprints corresponding to KLF10 (416 sites) and KLF11 (363 sites) in the TKO cortex (Figure 6H), indicating enhanced genomic occupancy of these regulators. In addition, the absence of detectable footprints at these sites in control cortices suggests *de novo* recruitment of KLF10 and KLF11 following the loss of H3K9me3 (Figure 6H). Collectively, these findings support a model in which H3K9me3 restricts TF access to TEs through chromatin compaction.

### H3K9me3 is associated with transcriptional repression of Cdkn1c

To identify genes potentially silenced by H3K9me3 that could directly contribute to the microcephaly phenotype observed in the TKO cortex, we focused on upregulated genes with known roles in suppressing cell proliferation. Among these, the imprinted gene *Cdkn1c* (also known as *p57Kip2*), which encodes a cyclin-dependent kinase inhibitor, emerged as a top candidate. *Cdkn1c* loss of function causes macrocephaly, whereas its upregulation suppresses progenitor proliferation and results in microcephaly^32–34^. However, the epigenetic mechanisms governing *Cdkn1c* expression in the developing brain remain poorly understood. Bulk RNA-seq revealed an approximately two-fold increase in *Cdkn1c* expression in the TKO cortex (Figure 7A). In control cortices, both the promoter and gene body of *Cdkn1c* were enriched for H3K9me3, whereas this mark was markedly depleted in the TKO cortex (Figure 7B). *Cdkn1c* expression in controls was primarily restricted to IPCs and immature cortical neurons, and *Cdkn1c* was selectively upregulated in these same populations in the TKO cortex (Figures 7C and 7D).

**Figure 7.**
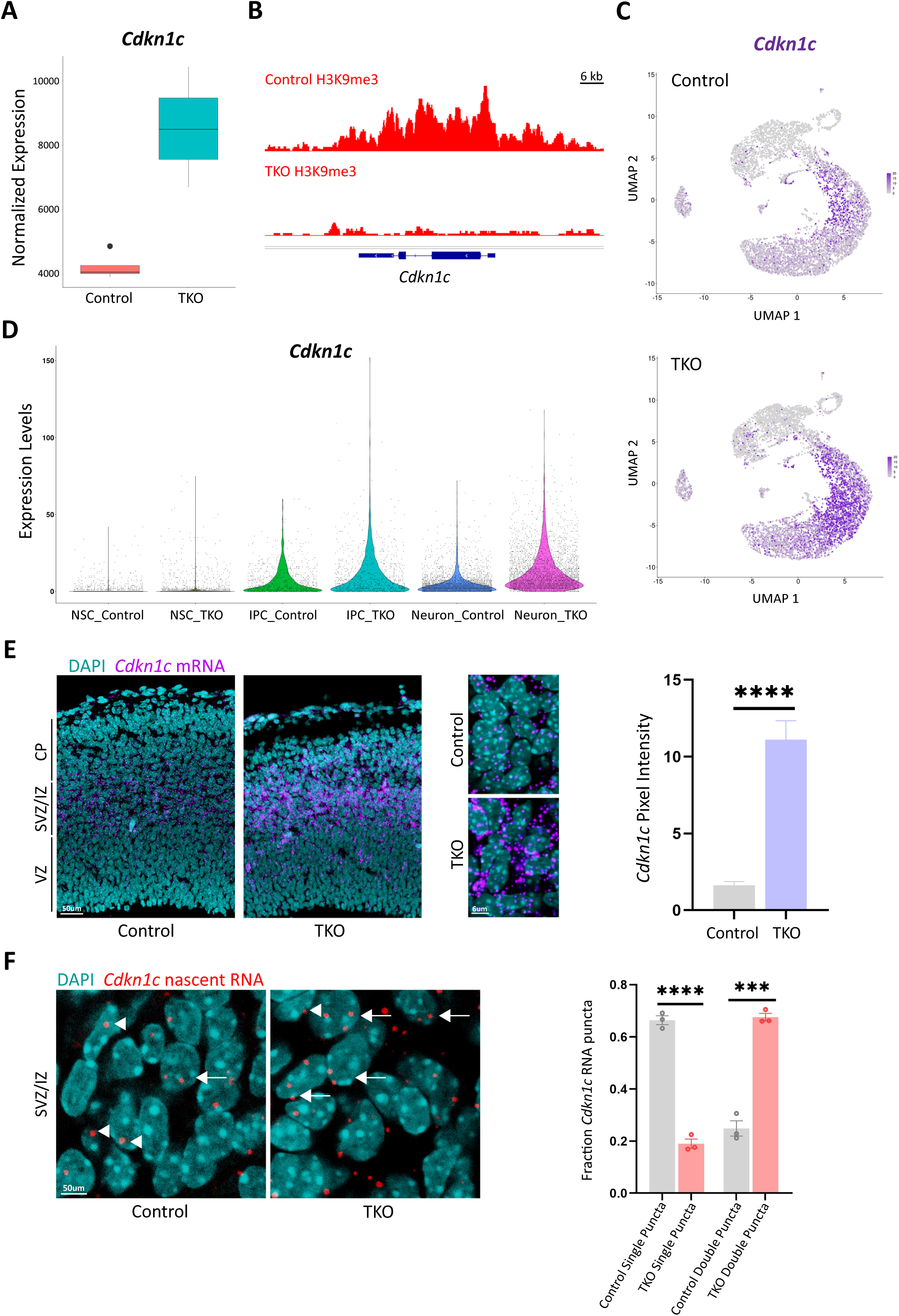
H3K9me3 is associated with *Cdkn1c* silencing in the developing cortex. (A) Bulk RNA-seq analysis revealed an approximately two-fold increase in *Cdkn1c* expression in the TKO cortex. (B) Genome browser tracks showing H3K9me3 enrichment across the promoter and gene body of *Cdkn1c* in the E14.5 control cortex. In contrast, H3K9me3 was markedly depleted across the *Cdkn1c* locus in the TKO cortex. (C) UMAP plot from scRNA-seq data showed *Cdkn1c* upregulation in the TKO cortex. (D) Violin plots from scRNA-seq analysis showed *Cdkn1c* expression in IPCs and neurons in the E14.5 control cortex, with significant *Cdkn1c* upregulation in these populations in the TKO cortex. (E) RNAscope *in situ* hybridization using a probe targeting *Cdkn1c* mRNA in E14.5 control and TKO cortices. Low-magnification images showed elevated *Cdkn1c* expression in the SVZ and IZ of the TKO cortex, indicating *Cdkn1c* expression in IPCs and immature neurons. Higher-magnification images highlight increased intensity and abundance of *Cdkn1c* mRNA puncta in TKO cells. (F) RNAscope *in situ* hybridization using a probe detecting nascent *Cdkn1c* RNA in E14.5 control and TKO cortices. Representative images from the SVZ and IZ showed that most nuclei in control samples displayed a single RNA punctum. In contrast, nuclei in TKO samples frequently exhibited two or more puncta, indicative of increased transcriptional activity. Arrowheads denote nuclei with a single punctum, and arrows indicate nuclei with two or more puncta. Quantifications are shown as mean ± SEM. Unpaired Student’s *t* test: \*\*\**p* < 0.001, \*\*\*\**p* < 0.0001.

RNAscope *in situ* hybridization in E14.5 control cortices confirmed high *Cdkn1c* mRNA levels in IPCs and immature neurons within the SVZ and IZ, respectively (Figure 7E). In addition, *Cdkn1c* mRNA levels were significantly increased in the TKO cortex (Figure 7E). To distinguish between enhanced mRNA stability and increased transcriptional activity, we performed RNAscope using a probe targeting nascent *Cdkn1c* transcripts. Nascent transcripts appeared as discrete nuclear puncta distinct from cytoplasmic mRNA puncta (Figure 7E and 7F). Whereas most control nuclei showed single *Cdkn1c* nascent RNA puncta, the majority of TKO nuclei exhibited two puncta (Figure 7F), suggesting enhanced gene transcription. Collectively, these findings support a role for H3K9me3 in silencing the growth inhibitory gene *Cdkn1c* in the developing cortex.

## DISCUSSION

This study identifies H3K9 methyltransferases as important regulators of cortical growth. Although we focused on the collective roles of SETDB1, SUV39H1, and SUV39H2 in cortical development, our data are consistent with a cooperative function for these enzymes, as reductions in *Setdb1*, *Suv39h1*, and *Suv39h2* dosage were associated with progressive cortical thinning. Our findings further suggest that the postnatal microcephaly phenotype reflects the combined effects of reduced progenitor proliferation, increased neuronal differentiation, and elevated cell death, although the relative contribution of each process remains unresolved. Notably, cortical thinning was detectable before the onset of apoptosis in TKO embryos, consistent with a role for altered cell cycle dynamics in the reduction of cortical thickness. Additionally, previous studies indicate that cell cycle lengthening shifts the balance from progenitor proliferation toward differentiation^35–38^. Therefore, our results suggest that cell cycle lengthening during mid-neurogenesis in the TKO cortex limits progenitor expansion and promotes premature differentiation at later stages, thereby contributing to microcephaly.

This work implicates H3K9me3 as a key regulator of chromatin condensation and transcriptional silencing during cortical neurogenesis. These conclusions are supported by several observations: (1) loss of H3K9me3 was associated with increased chromatin accessibility; (2) depletion of H3K9me3 at a subset of promoters correlated with transcriptional activation; and (3) loss of H3K9me3 was accompanied by enhanced transcription factor occupancy. These findings are consistent with *in vitro* studies demonstrating that combined silencing of *Setdb1*, *Suv39h1*, and *Suv39h2*, or mutation of histone H3K9, disrupts heterochromatin formation and transcriptional repression^15,28^. Despite these converging lines of evidence, establishing a direct causal relationship between H3K9me3 and repression of specific transcriptional targets will require further investigation. Future studies could include locus-specific restoration of H3K9me3 or conditional expression of catalytically inactive H3K9 methyltransferases^39,40^. In addition, loss of H3K9me3 in the embryonic cortex did not lead to widespread increases in chromatin accessibility, suggesting the presence of compensatory silencing mechanisms. Hence, the potential crosstalk between H3K9me3 and additional repressive pathways—such as DNA methylation and other histone modifications—warrants further exploration^41^.

H3K9me3 profiling during mouse and human embryonic development suggests that this mark plays a central role in establishing cell-type-specific transcriptional programs by silencing alternative lineage identities^16,42^. In line with this model, we observed that loss of H3K9me3 at a subset of promoters in the embryonic cortex was associated with aberrant activation of meiosis-related genes. These genes are normally restricted to germ cells and show minimal or no expression in the embryonic cortex. Consistent with our findings, depletion of H3K9me3 and concomitant upregulation of germ-cell related genes have also been reported in pluripotent stem cells *in vitro*^43^. Notably, the loss of other chromatin-modifying enzymes that promote heterochromatin formation during brain development results in ectopic activation of meiosis genes^44^. Examples include mutations in DNMT3B, MECP2, EHMT1/2, and KDM5C, all of which are associated with neurodevelopmental disorders^44^. Together, these observations indicate that partial activation of the germ cell transcriptional program may represent a shared molecular consequence of disrupted heterochromatin regulation^44^. Further studies will be required to determine the impact of ectopic meiosis gene expression on cell-cycle dynamics and neurogenesis in the developing brain, and to assess whether this phenomenon constitutes a common mechanism across multiple neurodevelopmental disorders.

Our results suggest that H3K9me3 is indispensable for silencing TEs in the developing cortex. Remarkably, a subset of TEs in the TKO cortex acquired features of active enhancers, including increased chromatin accessibility, H3K27ac enrichment, and transcriptional activity^45^. Consistent with these findings, recent studies have shown that loss of H3K9me3 in pluripotent and hematopoietic cells leads to activation of TEs that potentially function as cryptic enhancers^28,29^. Supporting this model, we found that activated TEs in the embryonic cortex were exclusively associated with upregulated genes in their genomic vicinity. Moreover, our data suggest that H3K9me3 restricts TF occupancy at a subset of promoters and TEs, likely through chromatin compaction. A similar role for H3K9 methylation has been described in invertebrates^46^. Finally, we observed enhanced occupancy of KLF family members at activated TEs, consistent with their reported binding to TE-derived enhancers in early embryos and pluripotent stem cells^47^. Together, these findings support a role for H3K9me3 in shaping the *cis*-regulatory landscape of the developing cortex by enforcing TE silencing.

Our data indicate that H3K9me3 contributes to repression of the cell cycle inhibitor *Cdkn1c* in the embryonic cortex. *Cdkn1c* is an imprinted gene whose expression is normally induced as NSCs transition to IPCs^32,48^. *Cdkn1c* upregulation leads to cell-cycle exit, depletion of the neural progenitor pool, and is associated with microcephaly^33,34^. In contrast, *Cdkn1c* loss results in enhanced progenitor proliferation and macrocephaly^32^. In the TKO cortex, increased *Cdkn1c* expression in IPCs coincided with reduced expression of cell cycle–associated genes, suggesting that elevated *Cdkn1c* promotes IPC differentiation and potentially contributes to the microcephaly phenotype. Additionally, H3K9me3 has been implicated in the regulation of genomic imprinting^49^. Hence, our findings also raise the possibility that loss of H3K9me3 leads to aberrant activation of the normally silenced paternal *Cdkn1c* allele. Consistent with this idea, overall *Cdkn1c* transcript levels were approximately doubled in the TKO cortex, and the average number of nascent *Cdkn1c* RNA puncta per nucleus was similarly increased. Alternatively, the elevated puncta number could also reflect a higher fraction of cycling cells containing sister chromatids due to altered cell-cycle dynamics. Discriminating between these possibilities will require allele-specific analyses capable of resolving maternal versus paternal *Cdkn1c* transcription and determining whether H3K9me3 more broadly safeguards genomic imprinting during corticogenesis.

### Limitations of this study

Our results suggest that H3K9me3 contributes to chromatin compaction and transcriptional repression in the embryonic cortex. However, it remains possible that non-catalytic functions of H3K9 methyltransferases, or methylation of non-histone substrates, also play important regulatory roles during neurogenesis. Dissecting these possibilities will require future studies employing catalytically inactive forms of H3K9 methyltransferases. More broadly, whether histone modifications are primary drivers of transcriptional regulation or arise as a consequence of gene expression changes remains an open question. Thus, H3K9me3 might act as a primary silencing mechanism at a subset of genes and regulatory elements, while reinforcing repression at others. Directly testing this model in specific tissues *in vivo* remains a significant technical challenge.

## MATERIAL AND METHODS

### Mouse strains

The conditional *Setdb1^lox/lox^* mouse line (*Setdb1^tm1.1Yshk^*) was crossed with a mouse line carrying a conditional *Suv39h1^lox/lox^* allele and a null *Suv39h2^KO/KO^* allele (*Suv39h2^em1Ksz^Suv39h1^em1Ksz^*/Mmjax, MMRRC strain #067176-JAX). The resulting conditional/null mouse line (*Setdb1^lox/lox^, Suv39h1^lox/lox^*, *Suv39h2^KO/KO^*) was then crossed with the *Emx1^Cre^* line (B6.129S2-*Emx1^tm1(cre)Krj^*/J, strain #005628-JAX) to produce TKO mice with cortex-specific deletion of *Setdb1* and *Suv39h1* over a null *Suv39h2* background. All TKO pups were runted and died within one day after weaning. Control animals were generated by crossing the conditional mouse line (*Setdb1^lox/lox^, Suv39h1^lox/lox^*, *Suv39h2^KO/KO^*) with C57BL/6 inbred mice (C57BL/6JOlaHsd; Inotiv) to generate mice harboring a functional copy of *Suv39h2* (*Setdb1^+/lox^, Suv39h1^+/lox^*, Suv39h2^+*/KO*^). Genotyping was conducted as previously described^16^. All animal housing, husbandry, pharmacological treatment, and euthanasia procedures were performed in accordance with a protocol approved by the Bloomington Institutional Animal Care and Use Committee (BIACUC) at Indiana University. The sex of mouse embryos and pups used in this study was not determined.

### Immunostaining

Embryonic forebrains were dissected and crosslinked overnight in 4% paraformaldehyde (PFA) at 4°C with mixing. For postnatal brain preparation, mouse pups were anesthetized with xylazine (20 mg/kg) and ketamine (150 mg/kg), and intracardially perfused with 1X PBS followed by 4% PFA. Brains were then dissected and post-fixed overnight in 4% PFA at 4°C with mixing. Crosslinked embryonic and post-natal brains were washed three times in 1X PBS, embedded in 4% agarose, and sectioned into 100 μm-thick slices using a VT1000 Vibratome (Leica Biosystems). Sections were incubated in antigen retrieval solution (10 mM sodium citrate, 0.05% Tween 20, pH 6.0) for 1 hr. at 70°C, followed by three washes in 1X PBS. Tissue sections were then blocked for 1 hr. at room temperature (RT) in blocking buffer (10% goat serum, 0.1% Triton X-100, 0.01% sodium azide in 1X PBS). Primary antibodies were diluted in blocking buffer and incubated with tissue sections overnight at 4°C with mixing. After three washes in 1X PBS, sections were incubated for 2 hrs. at RT with Alexa Fluor-conjugated secondary antibodies diluted in blocking buffer. Sections were washed three additional times in 1X PBS, and nuclei were stained with DAPI (4′,6-diamidino-2-phenylindole) before mounting sections on microscope slides using Fluoromount-G (Southern Biotech). We used the following primary antibodies: anti-H3K9me3 (Active Motif, 39062, 1:1000), anti-H3K9me2 (Abcam, ab1220, 1:1000), anti-Ki67 (Abcam, ab16667, 1:250), anti-CC3 (Cell Signaling Technology, 9661, 1:2000), anti-γH2A.X (Cell Signaling Technology, 9718T, 1:400), anti-PAX6 (Biolegend, 901301, 1:300), anti-TBR2 (Invitrogen, 14-4875-82, 1:250), anti-CUX1 (Santa Cruz, sc13024, 1:100), and anti-CTIP2 (Abcam, ab18465, 1:500). Secondary antibodies included anti-rabbit and anti-mouse IgG conjugated to Alexa Fluor 488, 546, or 647 (Invitrogen, 1:500). Images were acquired using a Leica SP8 confocal microscope and processed with LAS X Life Science microscope software (Leica Microsystems).

### EdU and BrdU labeling

EdU was prepared at 1 mg/ml in 1X PBS and administered to timed pregnant mice by intraperitoneal injection at a dose of 10 mg/kg body weight. EdU incorporation was detected in 100 μm vibratome sections using the Click-iT EdU kit with Alexa Fluor 647 dye (Invitrogen, C10340), following the manufacturer’s instructions. BrdU (Sigma-Aldrich, B5002) was prepared at 10 mg/ml in 1X PBS and administered to timed pregnant mice by intraperitoneal injection at a dose of 50 mg/kg body weight. BrdU incorporation was detected by immunostaining using anti-BrdU antibody (Abcam, ab6326, 1:200).

### RNAscope *in situ* hybridization

Dissected embryonic brains were crosslinked in 4% PFA, washed three times in 1X PBS, and cryoprotected by incubating overnight in 30% RNase-free sucrose (SIGMA, S0389) at 4°C with mixing. Cryoprotected tissues were embedded in O.C.T. embedding medium (Scigen), frozen, and sectioned at 10 μm using a cryostat. Sections were mounted on Superfrost Plus microscope slides (Fisher Scientific) and stored at - 80°C until use. RNAscope *in situ* hybridization was performed using the RNAscope Multiplex Fluorescent V2 Assay (ACDBio, 323100) following the manufacturer’s protocol with modifications. Briefly, cryosections were thawed at RT for 30 min., washed in 1X PBS for 5 min., and post-fixed in 4% PFA for 15 min. at 4°C. Sections were then washed in 1X PBS for 2 min. and treated with hydrogen peroxide for 10 min. at RT. RNAscope probes were hybridized for 2 hrs. at 40°C in a humidified chamber, followed by tyramide signal amplification and detection according to the manufacturer’s instructions. Probes used included a mouse *Cdkn1c* mRNA probe (Mm-Cdkn1c #458331) and a custom probe targeting nascent *Cdkn1c* RNA (Mm-Cdkn1c #1827271) that hybridizes to intronic sequences. The *Cdkn1c* mRNA probe was detected using Vivid 570 (1:1500), and nascent *Cdkn1c* RNA was detected using Vivid 650 (1:5000).

### RNA-seq

#### Bulk RNA-seq

Total RNA was extracted from E15.5 cortices using TRIzol reagent (Invitrogen, 15596026) and further purified with the RNeasy kit (QIAGEN, 74104). Four biological replicates were processed for both the control and TKO samples. RNA integrity was assessed using a TapeStation 2200 (Agilent). cDNA synthesis was conducted with the iScript cDNA Synthesis Kit (BIORAD, 1708890). RNA-seq libraries were prepared using the Illumina Stranded mRNA Library Prep Kit (Illumina, 20040534). Library concentration and size distribution were evaluated with a TapeStation 4200 (Agilent). Libraries were pooled and sequenced on an Illumina NextSeq 1000/2000 using a P2 v3 flow cell (100 cycles; catalog no. 20046811), configured to generate 2 × 59 bp paired end reads. Demultiplexing was performed using bcl2fastq2 Conversion Software v2.20.

#### Single-cell RNA-seq

E14.5 control and TKO cortices were dissected in Neurobasal-A minus phenol red (Gibco, 12349015) and pooled by genotype before dissociation. Tissues were incubated in 0.05% Trypsin-EDTA solution (Gibco, 25300062) plus 1% DNase (SIGMA, DN-25) for 10 min. at 37°C to generate single-cell suspensions. Following incubation, tissues were washed twice with 1 ml Hibernate E minus calcium (BrainBits, HECA500) and mechanically dissociated using a fire-polished Pasteur pipette in 1 ml of Hibernate E minus calcium containing 1% DNase I. Cells were centrifuged at 1000 rpm for 4 min. at 4°C and resuspended in DPBS without calcium and magnesium (Gibco, 14190144) supplemented with 0.04% BSA (SIGMA, A3059). Cell viability, assessed by Trypan Blue exclusion, ranged from 94–99%, and visual inspection confirmed the absence of doublets. Cells were diluted to ∼1200 cells/μl in 1 ml of DBPS plus 0.04% BSA. Single-cell libraries were prepared using the Chromium Next GEM Single Cell 3′ Reagent Kits v3.1 (10x Genomics, PN-100269) with the Dual Index Kit TT Set A (10x Genomics, PN-1000215), and sequenced on a NovaSeq X Series instrument using the 1.5B Reagent Kit (Illumina, 20104703).

### Native ChIP-seq

Embryonic cortices were stored at −80°C and thawed on ice for 5 min. Cortices were resuspended in 100 μl nuclear extraction buffer (10 mM Tris-HCl pH 8.0, 140 mM NaCl, 5 mM MgCl_2_, 0.6% NP40) and homogenized with a plastic pestle on ice until no tissue fragments remained. Samples were incubated on ice for 30 min., and nuclei were counted using Trypan Blue after the first 10 min. Approximately 2.3–3.6 million nuclei were used per ChIP-seq experiment. The micrococcal nuclease (MNase) master mix was prepared per ChIP sample by combining 15 μl 10X MNase buffer (NEB, M0247S), 2.2 μl dithiothreitol (DTT; 100 mM stock, Thermo Scientific, 707265ML), 1.5 μl halt protease inhibitor (100X stock, Thermo Scientific, 87785), 1.5 μl phenylmethylsulfonyl fluoride (PMSF; 100 mM stock, Thermo Scientific, 36978), 10 μl MNase (2 × 10^6^ U/ml stock, NEB, M0247S), and nuclease-free water to a final volume of 50 μl. For H3K27ac ChIP-seq, 3 μl sodium butyrate (2.5 M stock, SIGMA, B5887) were added. DNA digestion was performed by adding 50 μl of the MNase master mix to each sample and incubating for 12 min. at 37°C. The reaction was stopped by adding 1/10 volume of 100 mM EDTA, followed by 1/10 volume of 1% Triton X-100 / 1% sodium deoxycholate solution. Samples were incubated on ice for 15 min. and vortexed at medium speed for 30 s. Digested chromatin was supplemented with 1 ml of complete immunoprecipitation buffer (20 mM Tris-HCl pH 8.0, 2 mM EDTA, 150 mM NaCl, 0.1% Triton X-100, 1X halt protease inhibitor, 1X PMSF) plus 18 ng spike-in chromatin (Active Motif, 53083) and incubated for 1 hr. at 4°C. A 50 μl input control was collected before preclearing with 50 µl pre-washed Protein G magnetic beads (Invitrogen, 1003D) for 2-6 hrs. at 4°C with mixing. Precleared chromatin was then incubated overnight at 4°C with antibody-protein G bead complexes containing 5 μg anti-H3K9me3 (Active Motif, 39062) or anti-H3K27ac (Active Motif, 34034) antibodies and 2 μg of spike-in antibody (Active Motif, 61686) per ChIP sample. Beads were washed sequentially with 1 ml of complete immunoprecipitation buffer, two washes with low salt buffer (20 mM Tris-HCl pH 8.0, 2 mM EDTA, 150 mM NaCl, 1% Triton X-100, 0.1% SDS, 1X halt protease inhibitor), and two washes with high salt buffer (20 mM Tris-HCl pH 8.0, 2 mM EDTA, 500 mM NaCl, 1% Triton X-100, 0.1% SDS, 1X halt protease inhibitor). Chromatin was eluted in 200 μl of freshly prepared elution buffer (0.1 M NaHCO_3_, 1% SDS) and incubated for 2 hrs. at 65°C. Input controls were adjusted to 200 μl at this point. RNA was removed from samples by adding 4 μl 1 M Tris-HCl (pH 6.5), 4 μl 0.5 M EDTA, and 2 μl RNase A (stock at 20 μg/μl) and incubating for 1 hr. at 37°C. DNA was extracted in phase-lock tubes using 200 μl phenol: chloroform: isoamyl alcohol mix (25:24:1), vigorously mixed, and centrifuged at 15,000 *g* for 5 min. at 4°C. The aqueous phase was precipitated with 1/10 volume 3 M sodium acetate, 3 μl glycogen (20 mg/ml, Thermo Scientific, R0561), and 2.5 volumes absolute ethanol at −80°C for 2-3 hrs., followed by centrifugation at 15,000 *g* for 30 min. at 4°C. Pellets were washed with 70% ethanol and resuspended in DNA elution buffer (10 mM Tris-HCl pH 8.0). DNA was further purified using the DNA Clean & Concentrator Magbead Kit (Zymo Research, D4012). ChIP-seq libraries were prepared using the Ultra II DNA library Prep Kit for Illumina (NEB, E7103S) and NEBNext Multiplex Oligos for Illumina (NEB, E7335S). Sequencing was conducted as described in the bulk RNA-seq section.

### ATAC-seq

Aliquots of ∼50,000 cells were prepared as described in the scRNA-seq section. Each aliquot originated from individual E14.5 control and TKO cortices and was stored at −80°C until use. ATAC-seq libraries were generated using the Zymo-Seq ATAC Library Kit (Zymo, D5458) according to the manufacturer’s instructions. Library size distribution was assessed on a TapeStation 2200 (Agilent) before sequencing.

### Bioinformatics analyses

#### RNA-seq analysis

##### Bulk RNA-seq

Sequenced reads were processed for adapter removal and quality filtering using Trimmomatic v0.38^50^. Quality filtering applied a sliding window of three bases with a minimum average Phred quality score of 20, and reads shorter than 20 bases after trimming were discarded (Parameters: ILLUMINACLIP:adapters.fa:2:20:7 LEADING:20 TRAILING:20 SLIDINGWINDOW:3:20 MINLEN:20). Filtered paired-end reads were aligned to the mouse reference genome mm39 (GRCm39) using STAR v2.7.11a (Parameters: --outSAMattributes All --outFilterMultimapNmax 20 --seedSearchStartLmax 16 --twopassMode Basic)^51^. Read pairs that mapped uniquely and concordantly to exonic sequences of ENSEMBL-annotated genes (release 115) were quantified with featureCounts v2.1.1 (Parameters: -t exon -g gene_id -s 2 -p -B -C) from the Subread package^52^. Differential gene expression analysis was performed with DESeq2 v1.44.0^53^. For repeat element analysis, reads uniquely assigned to one or more copies of each repeat family were quantified, while reads mapping to multiple repeat categories were excluded. The resulting repeat count matrix was analyzed with DESeq2 v1.44.0 to identify differentially expressed repeat elements.

##### Single-cell RNA-seq

FASTQ files were processed using Cell Ranger v9.0.1 from 10X Genomics (https://www.10xgenomics.com/). The cellranger count pipeline was used to remove low-quality reads, align sequences to the mouse reference genome (mm10-2020-A), perform barcode calling, quantify unique molecular identifiers (UMIs), and produce gene-by-barcode expression matrices. Downstream analyses were conducted with Seurat v5.3.1^54^ using R v4.5.1. Quality control was performed by evaluating sample-specific metrics, including total UMI counts per barcode, number of detected features per barcode, and the proportion of mitochondrial reads. After QC filtering, data were normalized with SCTransform, integrated across samples, and subjected to principal component analysis (PCA) for dimensionality reduction. Cell clusters were visualized using uniform manifold approximation and projection (UMAP). Marker genes for each cluster were identified with the FindMarkers function, enabling annotation of NSCs, IPCs, and cortical neurons. Differential gene expression between controls and TKO cells within each cell population was assessed using the Wilcoxon test implemented in FindMarkers, applying an adjusted p-value threshold ≤ 0.05. Embryonic sex was inferred based on *Xist* expression across control and TKO barcodes. Barcodes with more than one *Xist*-specific UMI were classified as *Xist*^+^ (female), and all others as *Xist*^-^ (male). The TKO library originated from a pool of two male embryos, whereas the control library included one male and two female embryos. To mitigate sex-related transcriptional differences, genes differentially expressed between *Xist*^+^ and *Xist*^-^ control cells were identified using the same Seurat workflow and excluded from the final control versus TKO differential expression analyses. Gene ontology analysis of differentially expressed genes was conducted using ShinyGO 0.77^55^.

#### ATAC-seq analysis

Raw reads were trimmed using fastp v1.0.1 with the ‘--detect_adapter_for_pe -g’ parameters and aligned to the mm39 reference genome (BSgenome.Mmusculus.UCSC.mm39chrs.fa) using HISAT2 v2.2.1 with the ‘--no-spliced-alignment --no-unal --no-mixed --no-discordant’ parameters^56,57^. Duplicate reads were removed using MarkDuplicates from the Picard Toolkit v3.4.0 (https://broadinstitute.github.io/picard/, Broad Institute). Alignments were then shifted using alignmentSieve with ‘--ATACshift’ from deepTools v3.5.5^58^. Reads mapping to mitochondrial DNA (ChrM) or with a MAPQ score < 2 were removed using SAMtools v1.22.1^59^. Because embryonic sex was not determined, reads mapping to ‘ChrX’ and ‘ChrY’ were also excluded. bigWig files were generated using bamCoverage from deepTools v3.5.5 with the ‘--extendReads’ parameter. Peak calling was performed using MACS3 v3.0.3 with the parameters ‘--broad --shift −37 --extsize 73 -p 1e-2’^60^. Blacklisted genomic regions were compiled in ‘mm39.excluderanges.bed’ and removed from the BED files using the BEDTools v2.31.1 intersect command with the ‘-v’ argument^61,62^. Peak reproducibility was assessed using IDR v2.0.4.2 with ‘--rank p.value’ and a cutoff value of 540^63^. Differential accessibility analysis was performed using DiffBind v3.18.0^64^ executed in custom R scripts. ATAC-seq footprinting was carried out using the TOBIAS_snakemake pipeline^31^ on peaks showing increased accessibility in the TKO cortex. Peaks were annotated and motif analysis was conducted with findMotifsGenome.pl from the HOMER suite v5.1 with the parameter ‘-size −50,50’^65^.

#### ChIP-seq analysis

Analyses of H3K9me3 and H3K27ac ChIP-seq datasets were performed independently using the same workflow unless otherwise specified. All ChIP-seq experiments were normalized for immunoprecipitation efficiency using *Drosophila melanogaster* spike-in chromatin^66^. The normalized raw reads were trimmed using fastp v1.0.1 with the ‘--detect_adapter_for_pe -g’ parameters^56^ and aligned to a combined mouse and *Drosophila*reference genome (BSgenome.Mmusculus.UCSC.mm39chrs.fa and BSgenome.Dmelanogaster.UCSC.dm6chrs) using bowtie2 v2.5.4^67^. Mouse- and Drosophila-derived reads were separated using split_bam.py from the Spiker package v1.0.5^68^. Duplicate reads were removed using MarkDuplicates from the Picard Toolkit v3.4.0, and unpaired reads were excluded with reformat.sh from BBTools v37.62. Read counts for each ChIP sample were scaled by a factor equal to the minimum number of dm6-mapped reads across all samples divided by the number of dm6-mapped reads for that sample^66^. Peak calling was conducted with MACS3 v3.0.3 with a relaxed p-value cutoff (-p 1e-2)^60^. The ‘--broad’ parameter was applied for H3K9me3 ChIP-seq. Blacklisted genomic regions were compiled in ‘mm39.excluderanges.bed’ and removed using the BEDTools v2.31.1 intersect command with the ‘-v’ argument^61,62^. Peak reproducibility was determined using IDR v2.0.4.2 with ‘--rank p.value’ and a cutoff value of 540^63^. Differential enrichment analysis was performed using DiffBind v3.18.0^64^ executed in custom R scripts. For external H3K4me3 ChIP-seq datasets^69^, raw reads were trimmed using fastp v1.0.1 and aligned to the mm39 mouse reference genome using HISAT2 v2.2.1 with the parameters ‘--no-spliced-alignment --no-unal --no-mixed --no-discordant’. Duplicate reads were removed using MarkDuplicates. The heatmaps for all ChIP-seq datasets were generated from peaks called with MACS3 v3.0.3 using a relaxed p-value cutoff (-p 1e-2) followed by IDR analysis.

#### Identification of putative H3K9me3 target genes

To identify genes enriched for H3K9me3 at promoter regions, MACS3 v3.0.3 peak calling was performed with the default q-value threshold (-q 0.05). Reproducible peaks between biological replicates were identified using BEDtools v2.31.1 intersect with the parameters ‘-f 0.3 -r’. This approach provides a more comprehensive set of reproducible peaks than IDR, which can be biased toward weaker replicates^63^. Reproducible H3K9me3 peaks within ±1.5 kb of TSS were assigned to proximal promoter regions of putative target genes. These genes were subsequently intersected with the list of dysregulated genes in the bulk RNA-seq dataset to identify candidates potentially regulated by H3K9me3.

#### Binding and Expression Target Analysis (BETA)

To associate differentially expressed genes from bulk RNA-seq with distal ATAC-seq peaks showing increased accessibility in the TKO cortex, BETA-plus v1.0.7 was applied with the default 100 kb genomic window and the ‘-c .05’ parameter^30^. ATAC-seq peaks located within ±1.5 kb of annotated TSS were excluded to focus on distal regulatory elements.

#### Integration of ATAC-seq and ChIP-seq data

ChIP-seq reads mapping to ‘ChrX’, ‘ChrY’, and ‘ChrM’ were removed using SAMtools v1.22.1 for consistency with ATAC-seq analyses. Scaling factors were calculated from BAM files using the normFactors method in the CSAW v1.38.0 package with 10 kb genomic windows^70^. BAM files were converted to bigwig format using bamCoverage with ‘--binSize 1’, and replicate bigwig files were combined using bigwigAverage, both commands from deepTools v3.5.5. Enrichment matrices were generated using deepTools v3.5.5 computeMatrix using the parameters ‘--referencePoint center -a 5000 -b 5000--missingDataAsZero --skipZeros’^58^. Heatmaps and signal profile projections were generated using plotHeatmap and plotProfile from deepTools v3.5.5, respectively^58^.

### Statistical analysis

#### Image quantification

Quantification of RNAscope puncta was performed manually using the cell counter tool in ImageJ/Fiji (National Institutes of Health, USA). Orthogonal optical plane visualization was used to ensure that nascent RNA puncta were correctly assigned to the appropriate nucleus and not inadvertently attributed to adjacent nuclei. Pixel intensities of RNA puncta were normalized using the same approach described below for immunostaining analyses.

Quantification of histone methylation immunostainings was performed on single-plane confocal images converted to 16-bit format in ImageJ using the default pixel intensity threshold. Regions of interest (ROIs) were manually defined based on the DAPI channel overlaid with the immunostaining channel, and pixel intensities were measured in 30-50 ROIs per condition. To account for variability in signal intensity across experiments, pixel intensities were normalized using the formula: adjusted intensity = (X – X_min) / (X_max – X_min), where X represents the pixel intensity of the histone methylation signal within a nucleus, whereas X_min and X_max correspond to the minimum and maximum pixel intensities observed across control and TKO samples. Statistical significance was assessed using a two-tailed unpaired Student’s t-test in Prism (GraphPad) and reported as \**p* < 0.05, \*\**p* < 0.01, \*\*\**p* < 0.001, \*\*\*\**p* < 0.0001.

#### Quantification of microcephaly, cell number, and cell density

Cortical thickness was measured as the distance (in microns) from layer I to the white matter tracks using the LAS X Life Science microscope software (Leica Microsystems) and anatomically equivalent regions of the somatosensory cortex. Cell counts were obtained either manually using ImageJ/Fiji or with the LasX cell counter software. At least three brains per control and TKO condition, derived from a minimum of two litters, were analyzed. Cell density was calculated by dividing the total number of nuclei by a defined cortical area (200 µm width × measured cortical thickness), and values were converted to cells/mm² to allow direct comparison across samples.

#### Quantification of cell cycle dynamics

Cell cycle length was determined using the thymidine analogs EdU and BrdU in combination with Ki67 immunostaining^19^. Following a 30 min. EdU pulse, S-phase cells (S cells) were identified as EdU⁺, indicating active DNA synthesis. Cells exiting S-phase between the EdU and BrdU labeling interval (L cells) were identified as BrdU⁺ EdU⁻. The total proliferative population (P cells) was identified as Ki67⁺. S-phase duration (Ts) was calculated as: Ts = (S cells / L cells) × 1.5 hrs., where 1.5 hrs. represents the interval between EdU and BrdU administration^19^. Total cell cycle length (Tc) was estimated using the formula Tc = (P cells / S cells) x Ts^19^. The cell cycle exit index was calculated as the fraction of EdU⁺ Ki67⁻ cells relative to the total number of EdU⁺ cells approximately 24 hrs. after EdU injection. A minimum of three brains from at least two litters were analyzed per condition. Significance was determined using an unpaired Student’s t-test and reported as: \**p* < 0.05, \*\**p* < 0.01, \*\*\**p* < 0.001, \*\*\*\**p* < 0.0001.

## Acknowledgements

We are grateful to Dr. Kenneth Zaret (University of Pennsylvania) and Dr. Yoichi Shinkai (RIKEN) for generously providing the *Suv39h1^lox/lox^*, *Suv39h2^KO/KO^*, and *Setdb1^lox/lox^* mouse lines. We thank Douglas Rusch, David Merritt, Jun Yin, and Reza Abdi at the Center for Genomics and Bioinformatics (Indiana University Bloomington) for their valuable support and advice with next-generation sequencing experiments and bioinformatic analyses. This work was supported by grants from the National Science Foundation (no. 2523526) and the Brain Research Foundation awarded to J.-M.B.

## Author contributions

Conceptualization: J.-M.B.; Investigation: S.W., J.-M.B.; Methodology: S.W., J.-M.B.; Validation: S.W.; Data curation: S.W., C.H., R.P.; Formal analysis: S.W., C.H., R.P.; Visualization: S.W., C.H., R.P.; Funding acquisition: J.-M.B.; Project administration: J.-M.B.; Supervision: J.-M.B.; Writing – original draft: J.-M.B.; Writing – review & editing: S.W., C.H., R.P., J.-M.B.

## Declaration of interests

The authors declare no competing interests.

## Data availability

The raw sequencing data generated in this study have been deposited in the NCBI Sequence Read Archive (SRA) under accession number PRJNA1401117.

## Supplemental information

Document S1. Figures S1–S5 and legends.

Table S1. DEGs identified with bulk and scRNA-seq, related to Figure 3.

Table S2. Putative gene targets of H3K9me3-mediated silencing, related to Figure 5.

Table S3. Repeat elements marked with H3K9me3 in the embryonic cortex, related to Figure 6.

Table S4. BETA analysis results, related to Figure 6.

---

**Figure S1.**
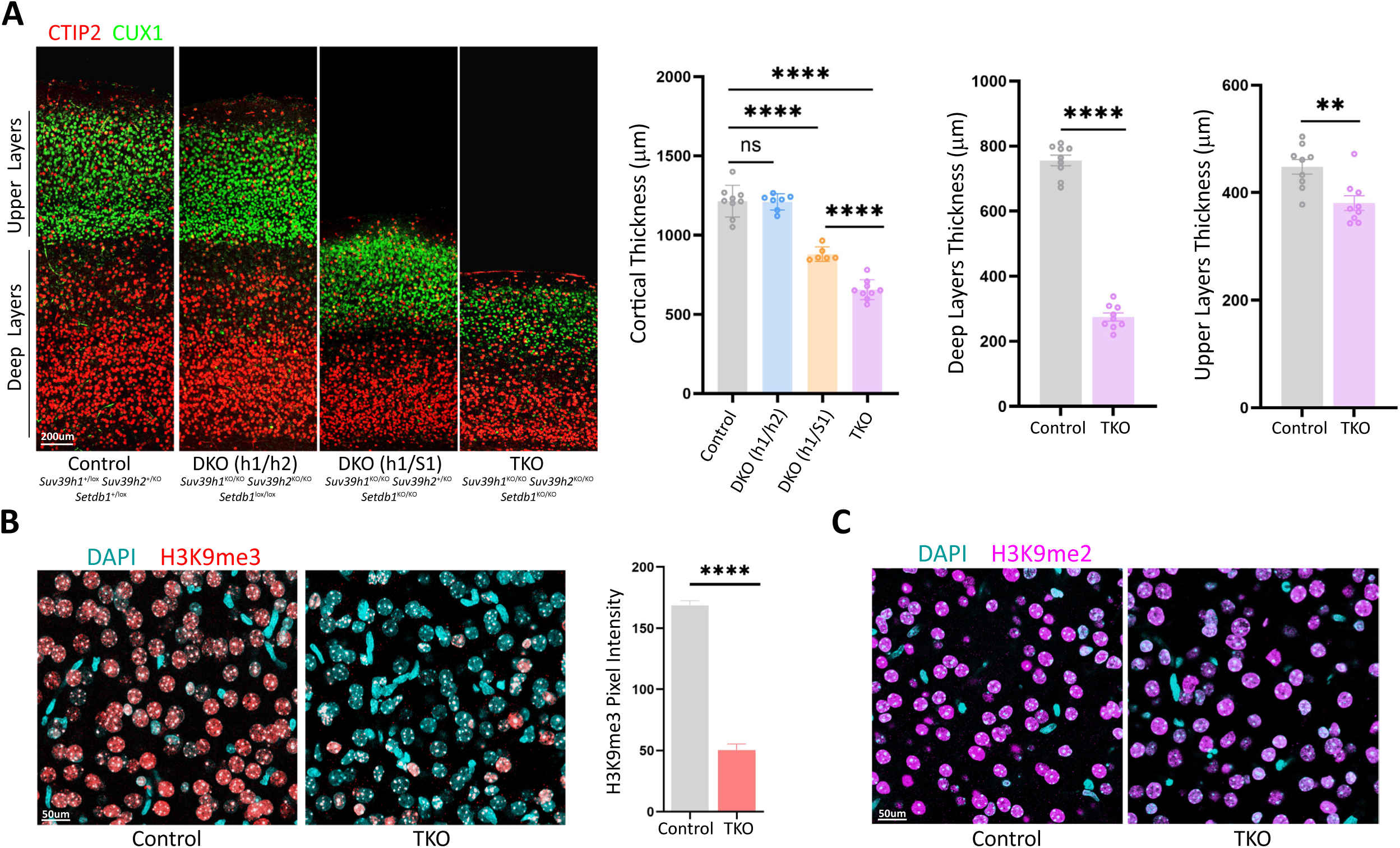
Cortical triple knockout of *Setdb1*, *Suv39h1*, and *Suv39h2* results in microcephaly and depletion of H3K9me3. (A) Coronal sections across P10 control, DKO, and TKO cortices stained for CUX1 (upper layers) and CTIP2 (deep layers). Complete genotypes are indicated for each sample. A significant reduction in cortical thickness was observed in the DKO (h1/S1) and TKO cortices, with the latter showing a more severe reduction. In the TKO cortex, the reduction in cortical thickness was more pronounced in deep layers than in upper layers. (B) Immunostaining revealed widespread depletion of H3K9me3 in the P10 TKO cortex. A small population of cells retained normal H3K9me3 levels, likely corresponding to cortical interneurons and ventrally derived glial cells, which are not targeted in this genetic model. (C) H3K9me2 levels did not show an apparent decrease in the P10 TKO cortex. Quantifications are shown as mean ± SEM. Unpaired Student’s *t* test: \*\**p* < 0.01, \*\*\*\**p* < 0.0001; ns, not significant.

**Figure S2.**
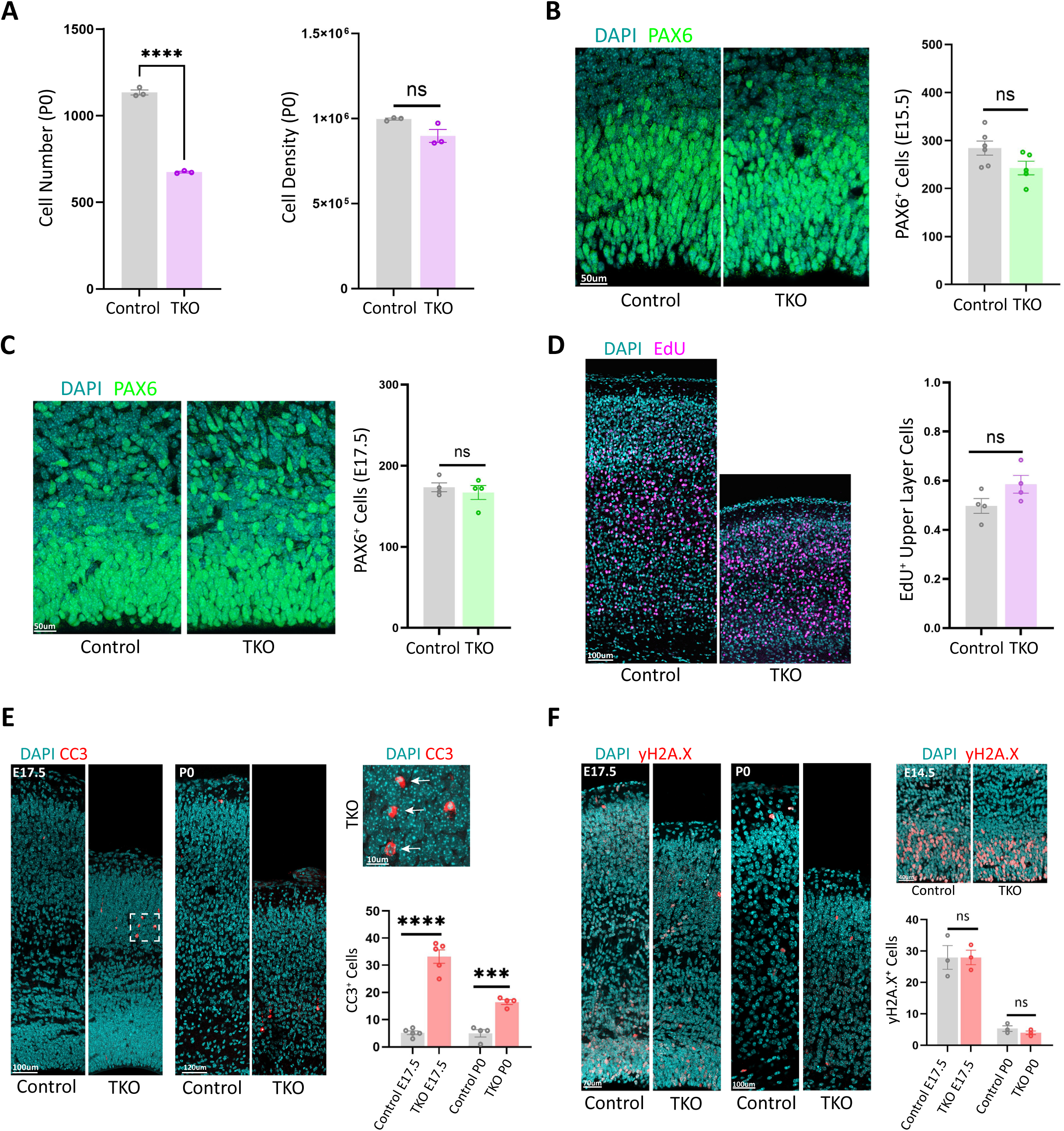
H3K9 methyltransferases regulate proliferation and neurogenesis in the embryonic cortex. (A) The total cell number within a cortical column was significantly reduced in P0 TKO cortices, whereas cell density was not significantly altered. (B) The number of PAX6⁺ cells did not differ significantly between E15.5. control and TKO cortices. (C) The number of PAX6⁺ cells did not change significantly in E17.5. TKO cortices. (D) EdU was administered at E13.5, followed by the analysis of EdU-labeled cells at P5 in control and TKO cortices. Although EdU labeling intensity was increased in TKO cells, the number of EdU⁺ cells in the upper cortical layers did not differ significantly between genotypes. (E) Quantification of cell death by CC3 immunostaining in E17.5 and P0 control and TKO cortices. The inset shows a higher-magnification view of CC3⁺ cells (arrows) in the E17.5 TKO cortex. (F) Quantification of genomic instability by γH2A.X immunostaining in E17.5 and P0 control and TKO cortices. No significant increase in γH2A.X⁺ cells was detected in the TKO cortex. Detection of γH2A.X in proliferating cells of E14.5 control and TKO cortices served as a positive control for the staining. Quantifications are shown as mean ± SEM. Unpaired Student’s *t* test: \*\*\**p* < 0.001, \*\*\*\**p* < 0.0001; ns, not significant.

**Figure S3.**
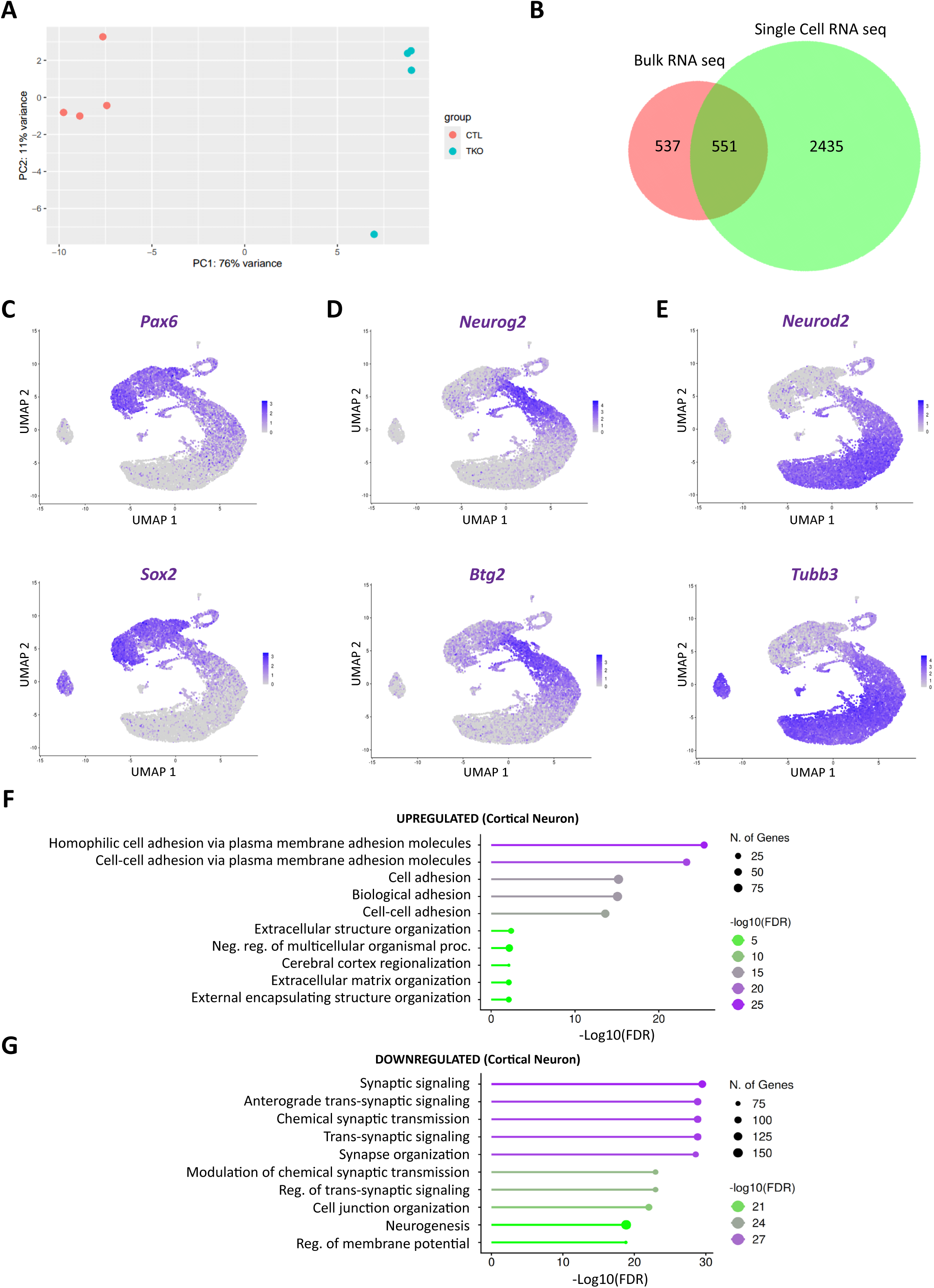
Bulk and single-cell transcriptomic analyses in control and TKO cortices. (A) Principal component analysis of bulk RNA-seq samples showing clear separation between control and TKO replicates. (B) Venn diagram illustrating the overlap of dysregulated genes identified in bulk and scRNA-seq datasets. (C) UMAP plots depicting high *Pax6* and *Sox2* expression in NSCs of the E14.5 control cortex. (D) UMAP plots showing elevated *Neurog2* and *Btg2* expression in IPCs of the E14.5 control cortex. (E) UMAP plots indicating *Neurod2* and *Tubb3* expression predominantly in projection neurons of the E14.5 control cortex. (F) Gene ontology analysis of genes upregulated in cortical neurons revealed enrichment for genes encoding cell adhesion molecules. (G) Gene ontology analysis of genes downregulated in cortical neurons revealed enrichment for genes involved in synaptic transmission.

**Figure S4.**
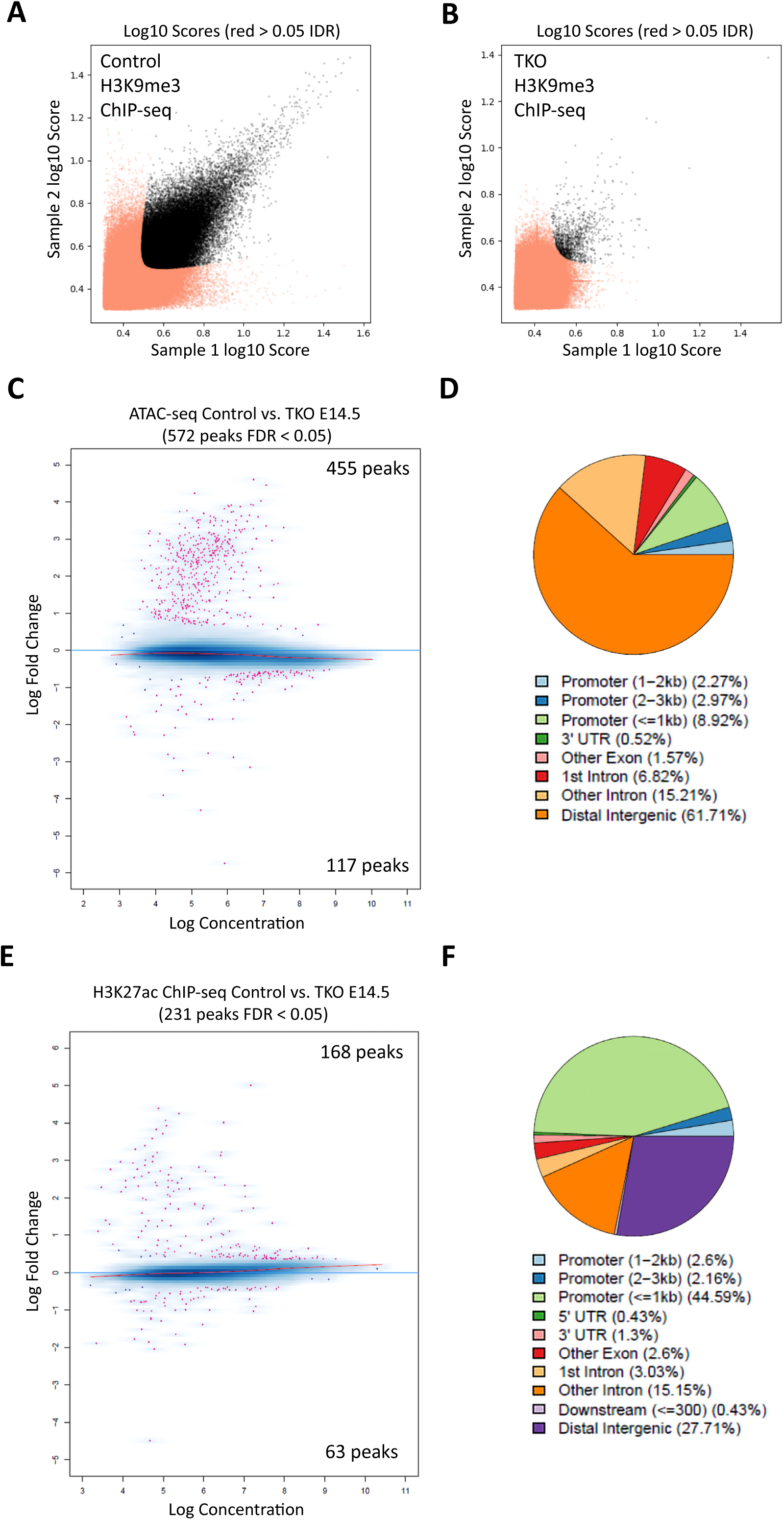
Chromatin analyses in control and TKO cortices. (A) IDR analysis of H3K9me3 ChIP-seq peaks in E14.5 control cortices comparing two replicates. Black dots indicate 51,053 reproducible peaks, and red dots represent non-reproducible peaks. (B) IDR analysis of H3K9me3 ChIP-seq peaks in two TKO replicates. Only 1,200 reproducible peaks were detected, consistent with H3K9me3 depletion in the TKO cortex. (C) Differential accessibility analysis in E14.5 control versus TKO cortices. The MA plot highlights 455 ATAC-seq peaks with increased accessibility and 117 peaks with decreased accessibility in TKO cortices (FDR < 0.05). (D) Genomic annotation of peaks with increased accessibility in TKO cortices shows enrichment in intergenic and intronic sequences. (E) Differential enrichment of H3K27ac ChIP-seq signals between E14.5 control and TKO cortices. A total of 168 regions showed increased H3K27ac, whereas 63 regions showed decreased H3K27ac in TKO cortices (FDR < 0.05). (F) Most regions with increased H3K27ac in TKO cortices were localized to promoters.

**Figure S5.**
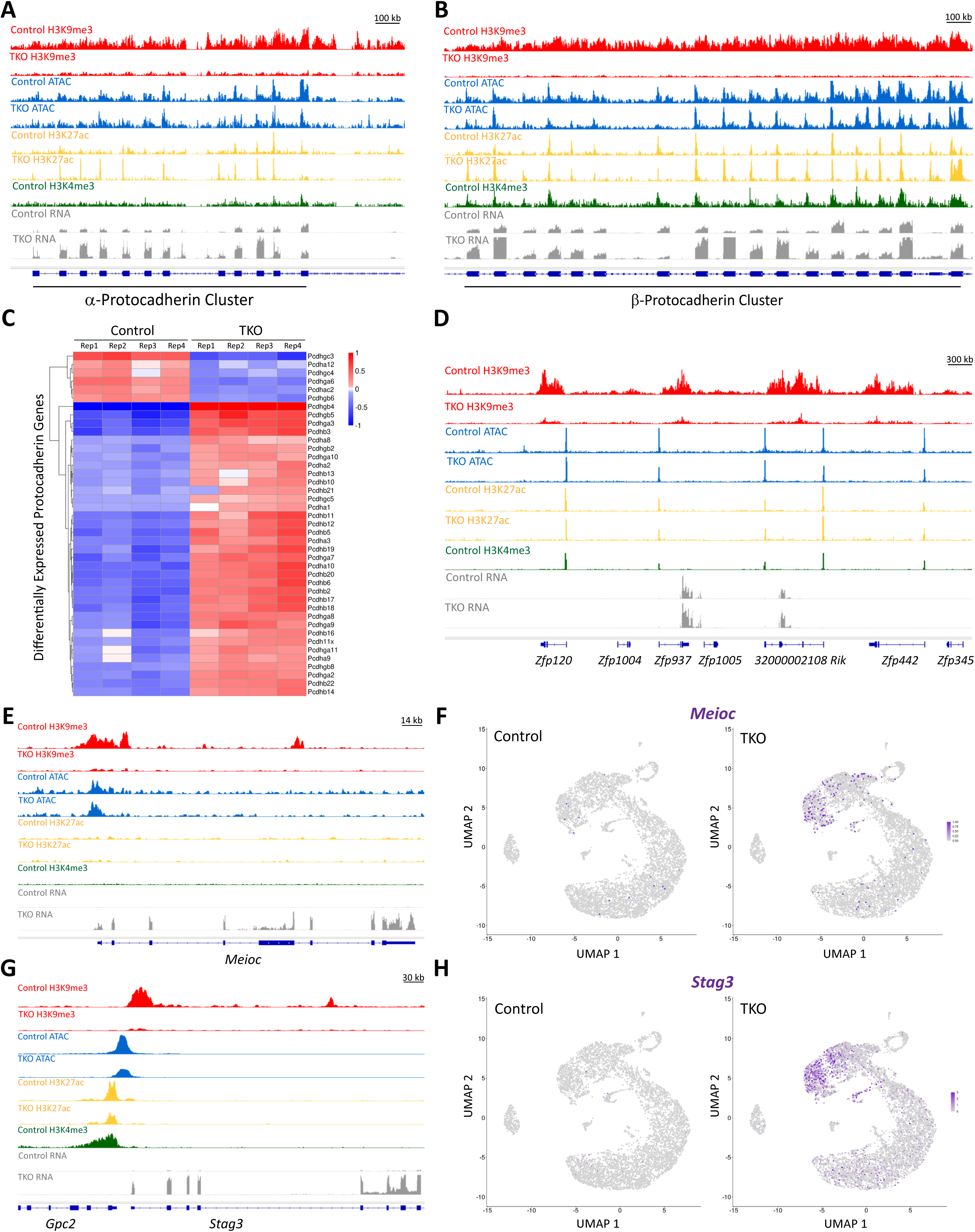
Loss of H3K9me3 is associated with upregulation of protocadherin clusters and meiosis genes in the embryonic cortex. (A) H3K9me3 decorated the α-protocadherin cluster in the E14.5 control cortex. Loss of H3K9me3 in the TKO cortex was accompanied by upregulation of protocadherin genes (RNA-seq tracks). (B) H3K9me3 decorated the β-protocadherin cluster in the E14.5 control cortex. H3K9me3 depletion in the TKO cortex was associated with upregulation of protocadherin genes (RNA-seq tracks). Chromatin accessibility within α- (A) and β-protocadherin clusters was largely unchanged between genotypes, whereas H3K27ac was locally increased in the TKO cortex. (C) Bulk RNA-seq analysis confirmed that most protocadherin genes were upregulated in TKO cortices. (D) Genome tracks showing H3K9me3 enrichment across a representative *Zfp* gene family in the E14.5 control cortex and H3K9me3 depletion in the TKO cortex. (E) H3K9me3 marked the promoter of *Meioc*, a meiosis-related gene activated upon H3K9me3 loss in TKO cortices. The low H3K4me3 signal at the *Meioc* promoter correlates with its minimal expression in control cortices. (F) scRNA-seq data showed that *Meioc* expression is primarily activated in NSCs of the E14.5 TKO cortex. (G) H3K9me3 decorated the promoter of *Stag3*, a meiosis-associated gene upregulated upon H3K9me3 depletion in TKO cortices. (H) scRNA-seq data show that *Stag3* expression is predominantly upregulated in NSCs of the E14.5 TKO cortex.

## Notes

### Competing Interest Statement

The authors have declared no competing interest.

